# A search for “perio-probiotics” by longitudinal dissection of oral bacterial community shifts during the onset and resolution of gingivitis

**DOI:** 10.1101/2022.08.25.505302

**Authors:** Michael W Hall, Nimali C Wellappuli, Ruo Chen Huang, Kay Wu, David K Lam, Michael Glogauer, Robert G Beiko, Dilani B Senadheera

**Affiliations:** Faculty of Graduate Studies, Dalhousie University, Halifax, Nova Scotia, Canada; University of Toronto, Toronto, Ontario, Canada; McMaster University, Hamilton, Ontario, Canada; University of the Pacific, San Francisco, California, USA; Faculty of Computer Science, Dalhousie University, Halifax, Nova Scotia, Canada; Stony Brook University, School of Dental Medicine, Stony Brook, New York, USA

**Keywords:** oral microbiome, gum inflammation, periodontal disease, gingivitis, 16S rRNA

## Abstract

**Aim:** To understand the spatiotemporal dynamics of bacterial succession during gingivitis, and to identify taxa with a critical role in gum health with prognostic value.

**Materials and methods:** Longitudinal microbiome data were collected from 15 individuals after completely discontinuing all forms of oral hygiene, and subsequently reintroducing it for three and two weeks, respectively. Sequences from the 16S rRNA V4-V5 gene region from sub- and supra-gingival plaque, saliva, and tongue sites were annotated and mapped to a reference tree of Human Oral Microbiome Database sequences.

**Results:** Suspending oral hygiene induced gingivitis, which was resolved after its resumption to baseline. Most significant shifts in bacterial abundance were observed in dental plaque, but not in saliva and tongue sites. During gingivitis-induction, baseline microbiota dominated by *Streptococcus*, was superseded by increased *Prevotella, Fusobacterium, Leptotrichia*, and *Porphyromonas* genera. Converse to its decline during disease-induction, gum health restoration was accompanied by a significant increase in streptococci.

**Conclusion:** We present the most comprehensive, spatiotemporal map of bacterial succession during gingivitis onset and resolution. We have identified taxa with potential as probiotic candidates for gum disease (i.e., perio-probiotics), and suggest tooth-associated plaque and not saliva or tongue plaque should be used in future prognostic tests.

## Introduction

Periodontitis is an infectious, inflammatory disease that is characterized by the destruction of tooth-supporting structures that include the alveolar bone, gingiva, and connective tissue (AlJehani, 2014;Nibali et al., 2013; Tumolo, 2013). The mildest form of periodontal disease that precedes periodontitis is known as gingivitis (Könönen et al., 2019). It is characterized by gingival inflammation in response to dental plaque accumulation, which can be restored to health with proper oral hygiene that reduces the bacterial burden. Left untreated, gingivitis can proceed to chronic periodontitis causing irreversible bone and tissue damage. Gingivitis - the early and reversible stage of the disease, therefore, represents a critical window to prevent further progression of disease and preclude permanent loss of bone and tissue. Hence, an understanding of transition of the microbiota from health to disease and vice versa, at an early stage can help identify useful prognostic targets and novel treatment modalities to halt or reverse disease progression before permanent damage occurs.

The pathogenesis of periodontitis is multifactorial, and involves complex interactions between host microbiota, immunity and environmental factors (e.g., smoking, diet) (AlJehani, 2014). The oral cavity contains heterogeneous niches, such as soft tissues of the tongue, saliva, and hard tooth surfaces, which harbour distinct microbiologically complex communities (Dewhirst et al., 2010; Wade, 2013;Human Microbiome Project Consortium and others, 2012;Ximénez-Fyvie et al., 2000). The causative role of plaque in the development of periodontal disease was first demonstrated using an experimental gingivitis model (EGM), by Löe *et al*. in 1965 (Löe et al., 1965). The non-invasive EGM comprises of two phases that allow the controlled induction and resolution of gingivitis in subjects with healthy gingiva. The induction phase is initiated by temporarily discontinuing all forms of oral hygiene practices (OHP) over several weeks. In the restoration phase that follows, gingival health is returned to baseline by resuming OHP, thus precluding permanent damage to the bones and dentition of study participants (Löe et al., 1965). EGM facilitates a non-invasive model in humans for longitudinal surveillance of oral microbiota (and other parameters) with disease progression (Löe et al., 1965;Offenbacher et al., 2009; Grant et al., 2010; Lee et al., 2012; Kistler et al., 2013; Huang et al., 2014).

Cumulative evidence from 16S rRNA gene and shotgun metagenomic sequencing studies suggest distinct bacterial community structures in health and periodontal disease (Abusleme et al., 2013; Galimanas et al., 2014; Tu et al., 2014; Szafranski et al., 2014; Lenartova et al.,2021; Marotz et al., 2022; Curtis et al., 2020). Recent studies conducted for metagenome and metatranscriptome analyses show that the disease-associated microbiota are distinct in their functional potential by significantly differentially expressing virulence-related functions that potentially promote tissue destruction and infection (Nowicki et al., 2018; Ai et al., 2017; Dabdoub et al., 2016). Despite these, only a limited number of human studies have been conducted focusing on temporal changes in the oral microbiome with gingivitis onset (e.g., (Huang et al., 2014, 2021; Grant et al., 2010; Lee et al., 2012; Kistler et al., 2013; Nowicki et al., 2018)), as opposed to more severe forms of periodontal disease. Further, most studies examining the early stages of disease have examined only the bacterial burden in a single location (e.g., subgingivae) or for a limited duration (Löe et al., 1965; Kistler et al., 2013; Grant et al., 2010; Lee et al., 2012; Huang et al., 2014; Nowicki et al., 2018). However, the oral cavity contains heterogeneous niches, such as tissues of the tongue, saliva, and hard tooth surfaces, which harbour distinct microbiologically complex communities (Dewhirst et al., 2010; Wade, 2013; Human Microbiome Project Consortium and others, 2012), which are yet to be characterized under longitudinal disease progression.

Taken together, the above limitations primarily involving cross-sectional studies have hampered our understanding how the bacterial consortia in varying oral sites shift when gums transition from healthy to inflamed ones for a prolonged time. In this study, we present the results of an EGM study that comprehensively characterizes how hundreds of bacterial taxa present in saliva, and plaque derived from subgingival and supragingival, as well as tongue dorsum sites change in community composition on a weekly basis in 15 individuals. In perhaps the most comprehensive and rigorous, longitudinal, 16S rRNA-based high-throughput microbiome study conducted to date, here we report oral microbiome data as participants develop gingivitis in a span of three weeks after stopping oral hygiene. What is also noteworthy about this work is that by restoring gum health by resuming oral hygiene in the last two weeks of the study, we present comprehensive maps of the bacterial communities when inflamed gums are transitioned from a diseased to a healthy state. Using this model, we report taxa that are particularly interesting because their relative abundance is initially decreased as gums are inflamed, and then significantly increased as gums become healthier at the end of the study. It is possible that these organisms can be promising candidates for probiotic therapy to treat or prevent gum disease. In addition, several taxa reported as significantly altered when gums transition from health to disease, and/or disease to health, can serve as prognostic bacterial candidate markers to track treatment-outcomes for gum disease. Importantly, our results show that only bacterial samples from tooth-associated plaque and not plaque derived from the tongue dorsum or saliva should be used in tests that track periodontal pathogens in disease-surveillance assays.

## Results

### Subject demographics, disease induction and resolution

Seven men and eight women aged 19 to 29 years were recruited for this microbiome study, which was conducted as a sub-study of a single-arm clinical trial that was originally designated to characterize oral and circulatory polymorphonuclear neutrophils during the onset and resolution of gingivitis (Wellappuli et al., 2017). All participants were systemically healthy and had no active or previous history of periodontal disease, chronic aphthous ulcers or tonsillitis.

During the 21-day induction period, analysis of all clinical parameters of gingival disease showed a progressive statistically significant increase (p < 0.01) in mean pocket depths, mean bleeding on probing, mean gingival index, and mean plaque index (Wellappuli et al.,2017) (Figure 1). All parameters of disease indices returned to baseline levels during the resolutions phase when normal oral hygiene was resumed (Wellappuli et al., 2017).

**Figure 1.**
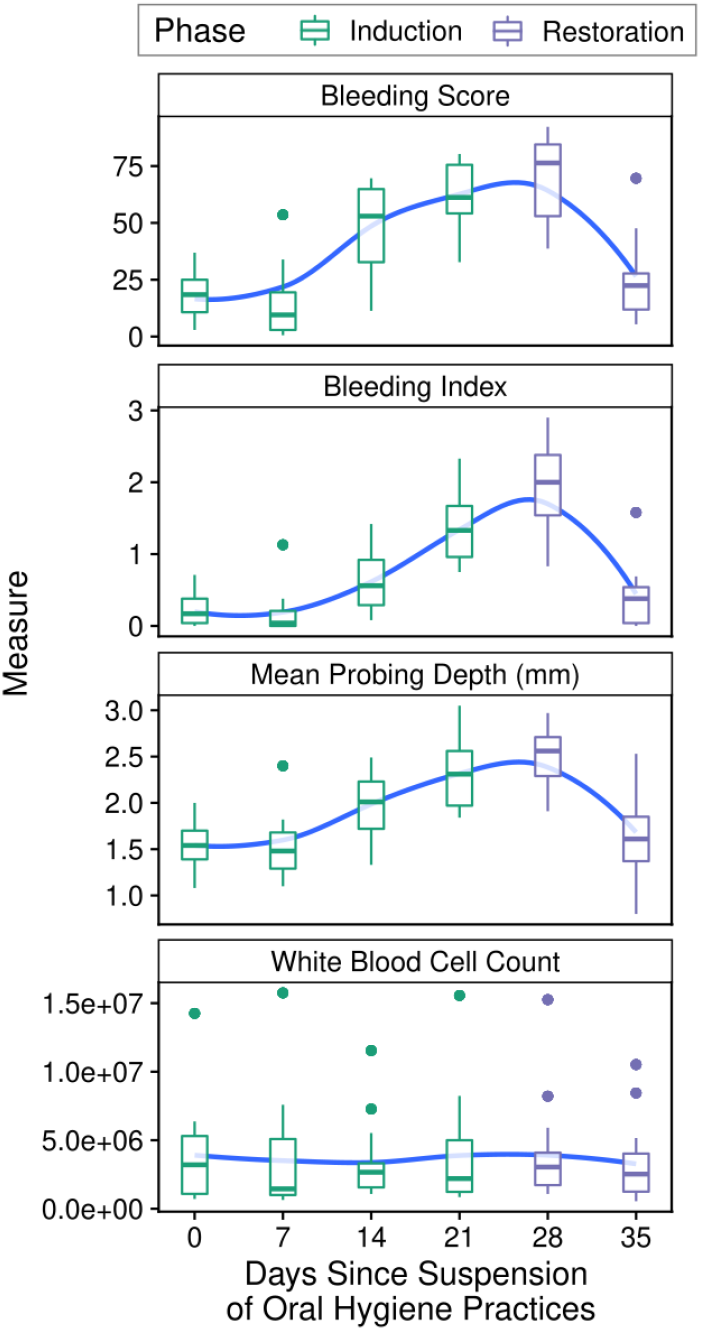
Indicators of periodontal health measured in subjects. Boxplots of indicators of periodontal health measured in Wellappuli et al. (2018), across time. Blue line represents a LOESS regression line.

### High-throughput sequencing of 16S rRNA gene

Post filtering and processing of 420 plaque and saliva samples, 12,899,647 16S rRNA gene sequences were obtained and clustered into 11,145 amplicon sequence variants (ASVs). Samples had a minimum of 4,222, maximum of 78,879, and median of 29,220 sequences. Rarefaction plots computed with the Shannon, Faith’s phylogenetic diversity, and observed ASV measures demonstrated a plateau in taxon accumulation (data not shown), suggesting samples were sufficiently deeply sequenced to capture the majority of the abundant taxa.

### Alpha diversity analysis

The alpha diversity, measured using the Shannon diversity and Observed ASVs measures, increased during the gingivitis induction phase in all four oral sites sampled (Figure 2). The magnitude of the slopes was largest, on average, in the subgingival site, followed by the supragingival site, saliva, and finally the tongue dorsum. Following the resumption of OHP, the alpha diversity levels returned to baseline levels during the restoration phase (Figure 2).

**Figure 2.**
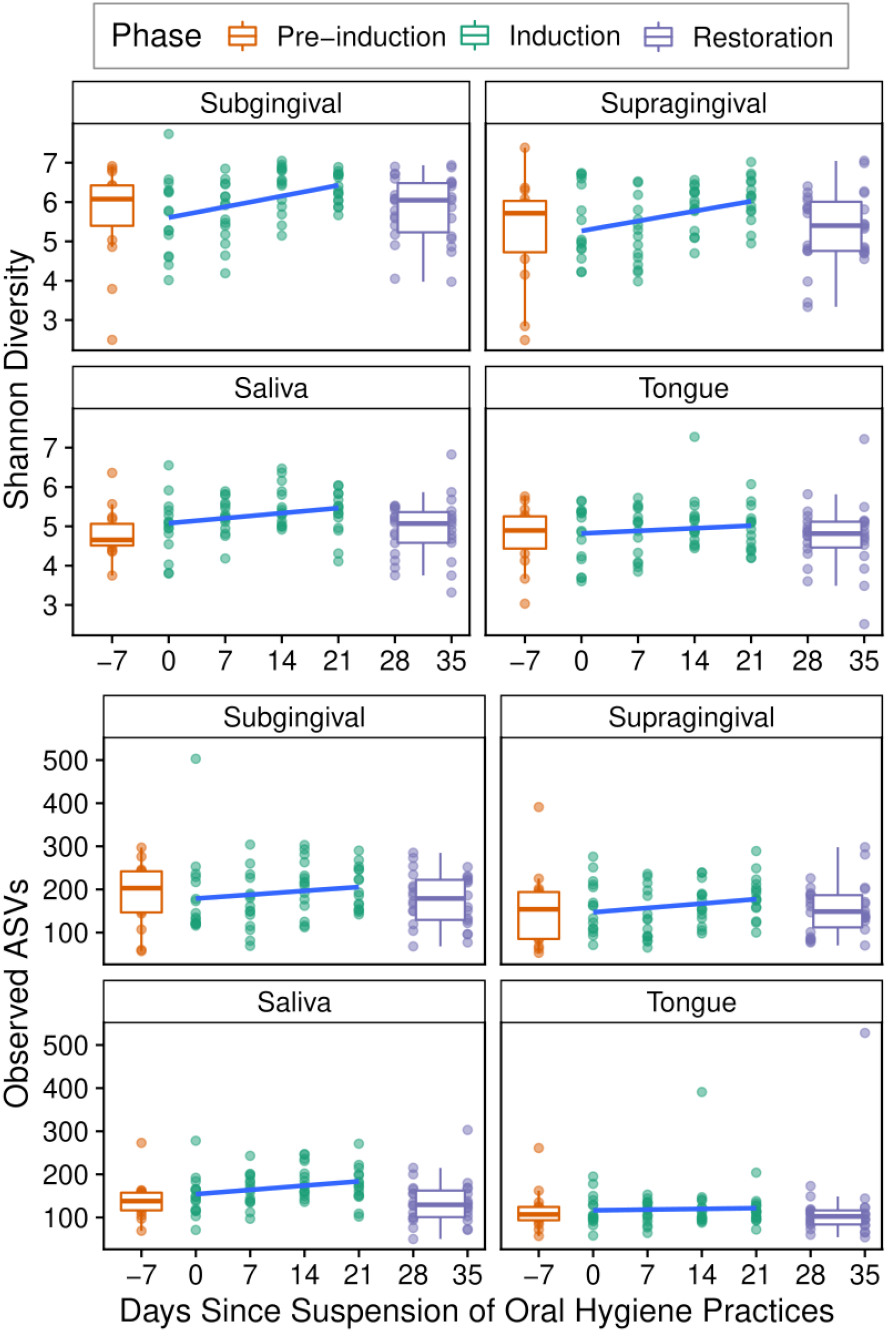
Alpha diversity over time in experimental gingivitis. Shannon diversity measures shown for each of the 420 samples. Boxplots show the median, 1st, and 3rd quartiles, and 1.5 times the inter-quartile range. Slope over induction phase (computed by GLM) shown in blue.

### Significant abundance trends in experimental gingivitis

All ASV representative sequences were taxonomically classified, with read counts aggregated at the phylum, genus, and species levels. Each taxonomic grouping had its CLR-transformed read counts tested for a non-zero slope over the induction phase. After p-value adjustment for multiple hypothesis test correction, 10 phyla, 40 genus-level designations, 43 species-level designations, and 66 ASVs were found to be significantly increasing or decreasing in relative proportion over the gingivitis induction phase. A graphical summary of the results at the phylum, genus, and ASV-level is shown in Figure 3 with full results in File S1.

**Figure 3.**
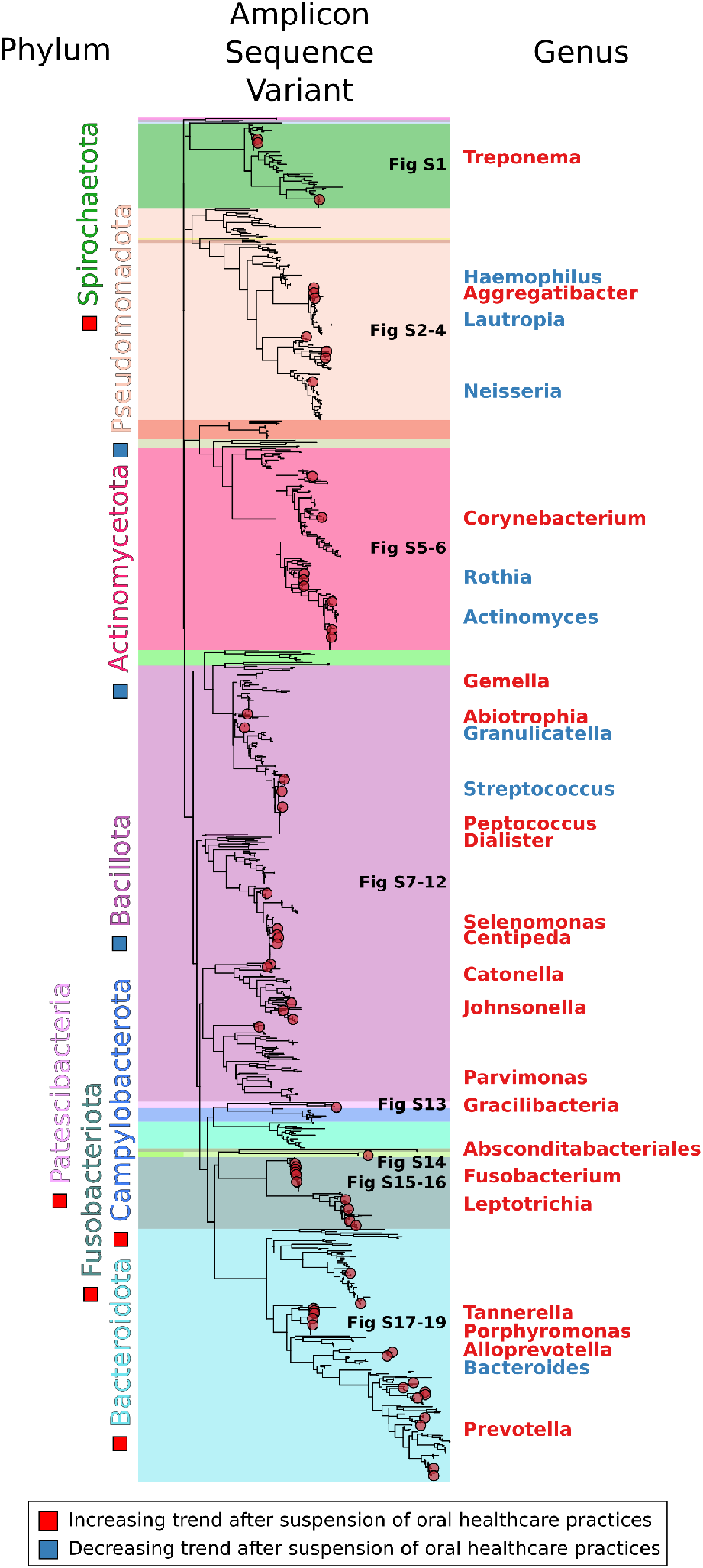
Summary of differentially abundant bacteria after suspension of oral health practices. A summary of the differentially abundant phyla, genera, and ASVs. Phylogenetic tree represents eHOMD V15.22 16S rRNA gene sequences trimmed to the V4-V5 region, with differential ASVs shown by red or blue circles. Regions of the tree are coloured by phylum-level classifications of the reference sequences. Red squares, circles, and genus labels indicate taxa that increased on aggregate over the gingivitis induction phase, with blue indicating a decreasing trend. Labels closer to the tree are the more abundant taxonomic groups.

At the phylum level, significant shifts in community proportions were observed in the subgingival and supragingival plaque. No statistically significant trends were observed at the phylum level in the tongue or salivary samples over the gingivitis induction period. From day 0 to day 21 after arresting OHP, the phylum-level dynamics were driven by increases in Bacteroidota (21.63% to 40.12% in subgingival and 18.78% to 32.78% in supragingival plaque) and Fusobacteriota (10.55% to 21.47% in subgingival and 9.96% to 18.86% in supragingival plaque) and decreases in Bacillota (34.63% to 24.48% in subgingival plaque), Pseudomonadota (21.29% to 8.42% in subgingival and 26.07% to 14.12% in supragingival plaque), and Actinomycetota (9.09% to 2.84% in subgingival and 10.30% to 5.24% in supragingival plaque). Several lower-abundance phyla (Spirochaetota, Campylobacterota, and Patescibacteria) also demonstrated significantly increasing trends, while two others (Desulfobacterota, and Cyanobacteria) demonstrated significantly decreasing trends in the tooth plaque samples (Figure 4, File S1).

**Figure 4.**
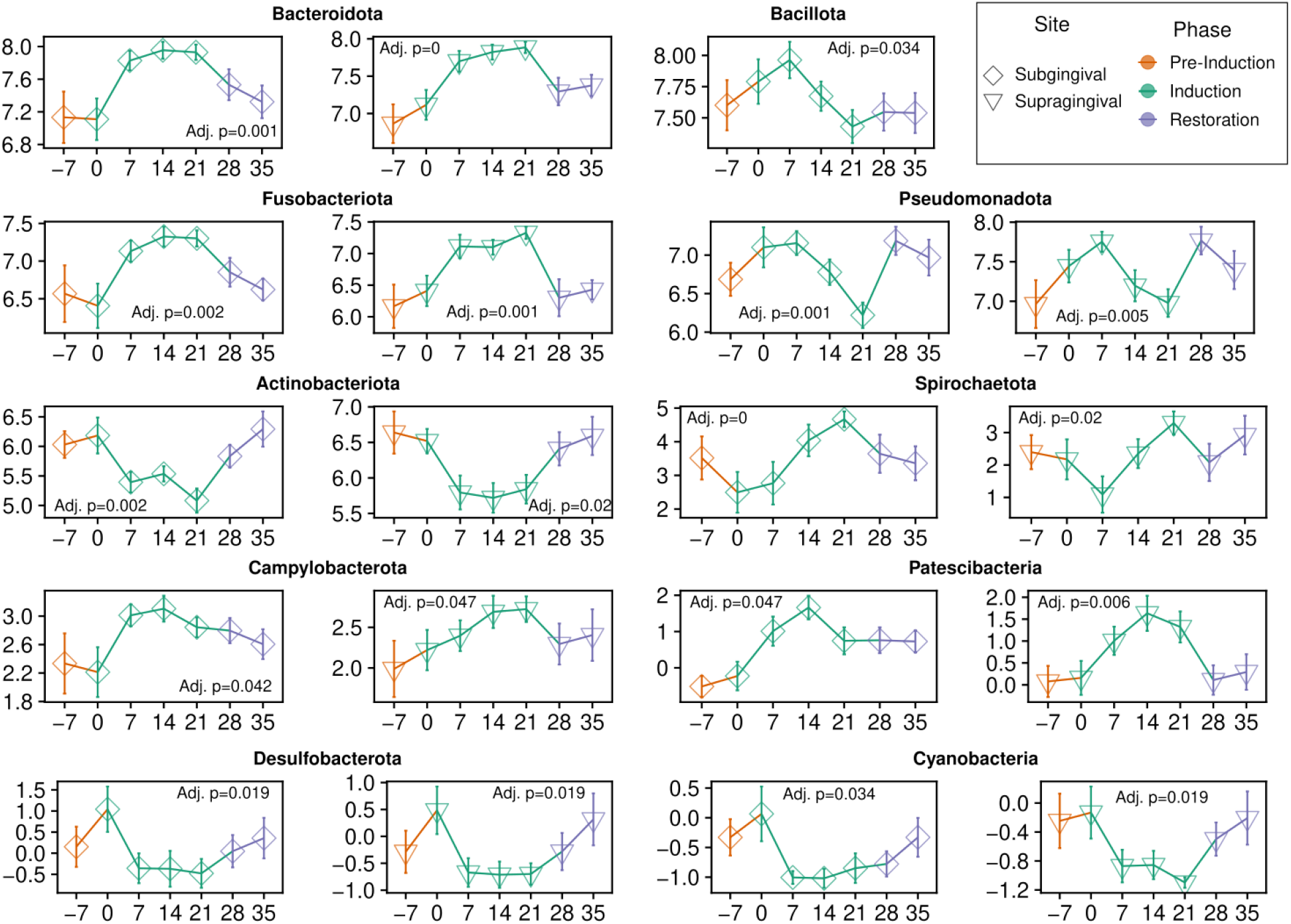
Differentially abundant phyla after the suspension of oral health practices. All phyla that were determined to be differentially abundant.

The most abundant genera with significantly increasing or decreasing community proportions are shown in Figure 5. Before the suspension of OHP, the most abundant genera across all subjects in the subgingival plaque samples were *Streptococcus* (17.89%), *Neisseria* (9.58%), *Prevotella* (8.92%), *Fusobacterium* (8.57%), and *Veillonella* (6.33%). After three weeks without OHP, the subgingival distribution shifted to *Prevotella* (21.00%), *Fusobacterium* (17.38%), Capnocytophaga (6.73%), *Veillonella* (4.85%), and *Porphyromonas* (4.54%).

**Figure 5.**
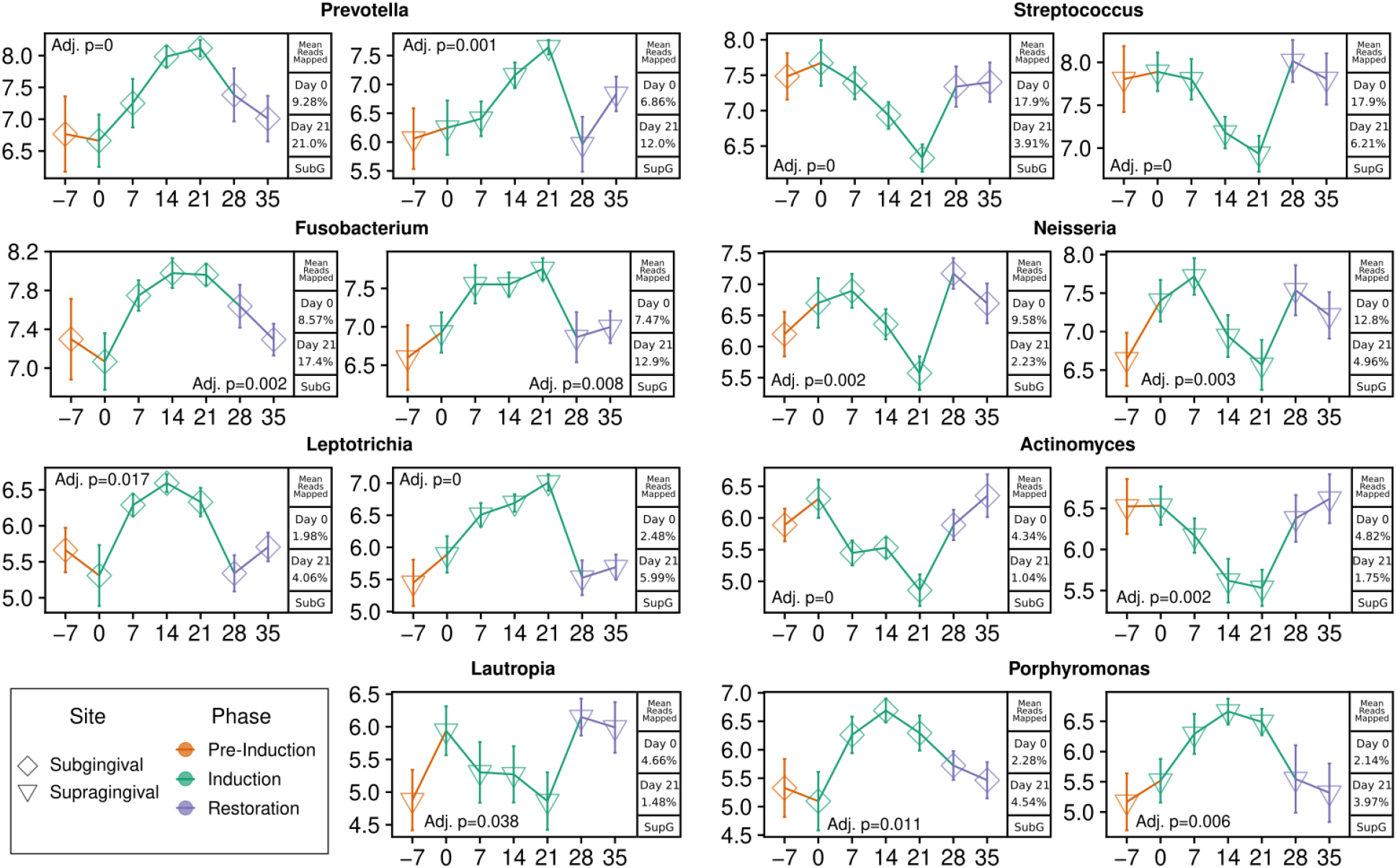
Top eight differentially abundant genera after the suspension of oral health practices.

Supragingival results mirrored this closely, with the most abundant genera at day 0 being Streptococcus (17.93%), *Neisseria* (12.82%), *Fusobacterium* (7.47%), *Prevotella* (6.84%), and *Veillonella* (6.83%). After gingivitis induction, that ranking changed to *Fusobacterium* (12.87%), *Prevotella* (12.17%), *Veillonella* (10.58%), *Capnocytophaga* (10.48%), *Leptotrichia* (5.99%), and *Neisseria* (4.96%).

The salivary genus composition was considerably more stable, with the most abundant genera at day 0 being *Veillonella* (22.71%), *Prevotella* (21.04%), *Neisseria* (10.51%), *Streptococcus* (10.33%), and *Fusobacterium* (7.75%). By day 21, the rank-order of genera remained the same at *Veillonella* (22.48%), *Prevotella* (18.14%), *Neisseria* (10.09%), *Streptococcus* (10.26%), and *Fusobacterium* (7.98%).

The genus-level composition of the tongue was similarly stable, with most abundant genera at day 0: *Prevotella* (20.17%), *Veillonella* (17.99%), *Streptococcus* (10.95%), *Neisseria* (10.05%), and *Fusobacterium* (8.35%). By day 21 the ranking changed to *Prevotella* (23.15%), *Veillonella* (17.83%), *Fusobacterium* (11.05%), *Streptococcus* (7.64%), and *Neisseria* (6.59%). In the tongue samples, exactly one CLR-transformed result was statistically significant: an increase of the genus *Aggregatibacter* from 0.01% to 0.09% mean relative abundance over the induction phase, representing a 6.6-fold increase.

A brief set of interesting bacterial dynamics after the suspension of OHP includes: the initial increase of *Neisseria* for the first two weeks after suspension of OHP followed by a rapid decline, the collapse of the genera *Haemophilus*, decreasing 30-fold from 3.41% to 0.11% in subgingival plaque, *Rothia*, decreasing 20-fold from 1.24% to 0.06% on average in subgingival plaque, and a similar 12-fold subgingival decrease in *Corynebacterium durum* from a mean of 0.45% to 0.04%, as well as a nearly 5-fold subgingival increase in *Alloprevotella* and 8-fold increase in *A. tannerae* specifically, and a statistically significant increase in *Cardiobacterium valvarum* in both tooth plaque environments (File S1).

### Socransky’s subgingival complexes after suspension of oral health practices

Given their historically cited role in periodontal disease, it is important to highlight the dynamics of the members of Socransky’s subgingival complexes after suspension of OHP (Socransky et al., 1998). Table 1 shows the taxonomic designations of Socransky’s complex (using taxonomic names from SILVA 138), the number of ASVs found in that group, and the subgingival abundance at the start (day 0) and end (day 21) of the experimental gingivitis induction phase. No sequences classifying to members of the purple complex were observed (Table 1). The green complex had mixed dynamics within its members, with *C. gingivalis* decreasing significantly and related species *C. sputigena* and *C. ochracea* roughly doubling in relative abundance, though these decreases did not result in a statistically significant slope across all induction time points.

**Table 1.** Members of Socransky’s complexes after three weeks of oral hygiene practice suspension. * indicates that this taxonomic group’s abundance increase/decrease was statistically significant (non-zero slope) at p≤0.05 after CLR transformation.

| Complex | Taxonomic Group | Number of ASVs | Subgingival Mean Reads (%) |  |
| --- | --- | --- | --- | --- |
|  |  |  | Day 0 | Day 21 |
| Red | *Treponema denticola* | 4 | 0.205 | 0.191 |
| Red | *Tannerella forsythia* | 3 | 0.095 | 0.061 |
| Red | *Porphyromonas gingivalis* | 2 | 0.005 | 0.112 |
| Orange | *Fusobacterium* * | 471 | 8.565 | 17.383 |
| Orange | *Prevotella intermedia* | 12 | 1.453 | 1.775 |
| Orange | *Prevotella nigrescens* * | 44 | 0.875 | 2.988 |
| Orange | *Campylobacter gracilis* | 3 | 0.084 | 0.098 |
| Orange | *Campylobacter showae* | 6 | 0.058 | 0.144 |
| Orange | *Eubacterium nodatum* | 2 | 0.010 | 0.003 |
| Orange | *Campylobacter rectus* | 0 | 0.000 | 0.000 |
| Orange | *Streptococcus constellatus* | 0 | 0.000 | 0.000 |
| Green | *Capnocytophaga gingivalis* * | 13 | 0.849 | 0.335 |
| Green | *Capnocytophaga sputigena* | 6 | 0.681 | 1.356 |
| Green | *Capnocytophaga ochracea* | 8 | 0.023 | 0.046 |
| Green | *Campylobacter concisus* | 7 | 0.011 | 0.014 |
| Green | *Aggregatibacter actinomycetemcomitans* | 0 | 0.000 | 0.000 |
| Purple | *Veillonella parvula* | 0 | 0.000 | 0.000 |
| Purple | *Schaalia odontolyticus* | 0 | 0.000 | 0.000 |
| Purple | *Selenomonas noxia* | 0 | 0.000 | 0.000 |
| Yellow | *Streptococcus* * | 356 | 17.886 | 3.908 |
| Blue | *Actinomyces* * | 226 | 4.341 | 1.041 |

The members of the red complex demonstrated mixed dynamics in early gingivitis onset. While none of the shifts were statistically significant across the four-week gingivitis induction period, *T. denticola* saw a small decrease in mean subgingival abundance between day 0 and 21 and *T. forsythia* saw its abundance decrease by a third, while *P. gingivalis* saw a large 22-fold increase, while still remaining very low abundance as a proportion of the overall community.

The yellow and blue complexes both saw significant decreases in their relative community importance during gingivitis induction. The yellow complex is populated by members of *Streptococcus*, which had three significantly decreasing ASVs found in three distinct clades for this gene region (Figure S8). The blue complex, which is comprised of *Actinomyces*, also had three significantly decreasing ASVs: two belonged to a clade containing the species *A. naeslundii, A. oris*, and *A. viscosus* and the third significant ASV belonging to a clade with *A. gerencseriae* (Figure S6).

The orange complex showed the most consistent dynamics, with all but one member (*Eubacterium nodatum*) seeing increases. However, only *Fusobacterium* and *Prevotella nigrescens* had statistically significant increases. Although no ASVs were classified as *C. rectus*, the gene region tree reveals little to no nucleotide differentiation between *C. rectus* and *C. showae* in the V4-V5 region (Figure S13). No ASVs were classified down to *Streptococcus constellatus*, and none of the significantly increasing or decreasing ASVs placed near the *S. constellatus* reference sequence within the *Streptococcus* reference tree (Figure S8).

## Discussion

Here we report the first EGM study to utilize high-throughput 16S rRNA gene sequencing to comprehensively investigate the transition of the oral microbiome in subgingival, supragingival, tongue dorsal sites and saliva during development of gingivitis and its resolution. This study was conducted as an extension of a single-arm clinical trial that characterized oral and circulatory polymorphonuclear neutrophils during the induction and resolution of experimental gingivitis (Wellap-puli et al., 2017). The EGM constituted a three-week gingivitis induction phase during which 15 study participants discontinued all forms of oral hygiene (i.e., brushing, flossing, etc.) that was followed by a final recovery phase, which included OHP resumption for two more weeks. By comparing data with that obtained from a baseline consortium derived before starting the induction phase, this study represents one of the highest resolution investigations conducted to date, during gingivitis induction. Overall, we identified trends in broad taxonomic groups (Figures 4, 5) as well as trends in specific amplicon sequence variants (Figure 3, Figures S1–S19) during the early onset of, and recovery from, gingivitis. In particular, disease onset was characterized by an increase in alpha diversity in the subgingival and supragingival plaques. After OHP were resumed, alpha diversity values and relative abundance of the majority of taxonomic groups moved back to the baseline, in step with the immunological and periodontal measures published in Wellappulli *et al*. (Figures 1, 4, 5, S20).

### Orange, yellow, blue complexes most dynamic subgingival complexes after three weeks without OHP

The microbial complexes described by Socransky *et al*. are reported to be major contributors to periodontal health and disease. As gingivitis can be a precursor to periodontitis, it is valuable to identify the dynamics of the microorganisms that comprise these complexes during the induction of gingivitis. The major stories within the subgingival complexes are the collapse of the yellow and blue complexes and increase in orange complex members.

In the red complex, while *P. gingivalis* appears to be rapidly gaining importance in the community from a very low abundance state in the early weeks of gingivitis (though statistically non-significant after correction), the other two members of the complex, *T. denticola* and *T. forsythia*, saw decreases in their mean abundance (Table 1).

In the orange complex, an increase of the diverse genus *Fusobacterium* and the species *P. nigrescens* are the only two statistically significant results. Although no sequences were annotated as *C. rectus* the reference gene tree shows that *C. showae* and *C. rectus* have little nucleotide variation in the V4-V5 hypervariable gene regions (Figure S13). It is possible that *C. rectus* is present and it is being misclassified as *C. showae* under our experimental parameters. On the other hand, no significant species or ASVs were detected near *S. constellatus* on the reference *Streptococcus* tree (Figure S8).

From the Streptococal tree, it was apparent that the yellow complex had four groups that decreased on aggregate: *S. mitis/oralis/infantis*, *S. sanguinis*, *S. parasanguinis/australis*, and *S. salivarius/vestibularis/thermophilus* (Figure S8). Similarly for the blue complex, *Actinomyces* had three ASVs in two species groups that decreased on aggregate: two ASVs in a group containing *A. oris/naeslundii/viscosus/johnsonii* and one ASV in a group with *A. gerencseriae/israelii* (Figure S6).

ASVs within the blue (*Actinomcyes*) and yellow (*Streptococcus*) complexes did not frequently classify down to species level. The phylogenetic trees (Figures S6 and S8)) help visualize why: clades within these groups often have multiple species designations with very little nucleotide variation between them. This results in an unacceptably low confidence score as it fails to differentiate between species with nearly identical V4-V5 gene regions, and the classification process defaults to a genus level annotation. Nevertheless, the phylogenetic reference trees help situate these ASVs relative to others and to reference sequences, and helps increase the resolution for those taxa. Future studies probing these microbial groups with DNA sequencing should consider a gene fragment or gene with higher within-group nucleotide diversity to clarify the uncertain phylogenies and classifications provided here and in studies with similar sequencing protocols.

### Restarting oral hygiene practices returns community to baseline

Our results show that the reestablishment of OHP after an extended period of suspension allows the bacterial community to revert to a state similar to that observed in the baseline. Alpha diversity values increase over time during induction of gingivitis, but decrease to baseline level after a “dental cleaning” procedure and resumption of daily OHP. The majority of taxonomic groups that were disrupted return to near the baseline levels observed after the restoration phase (time point −7, preinduction phase). The communities return to baseline (as shown by a-return to pre-induction phase abundance ranges) largely by time point 28 (e.g. Figures 4), and as the healthy community is re-established, subjects’ bleeding index, bleeding score, and mean pocket depth gradually return to their baseline one week later (time point 35) (Figure 1) (Wellappuli et al., 2017).

### Next steps in the search for perio-probiotics and diagnostic markers

In this work, we have identified broad microbial taxonomic groups and high-resolution sequence variants that shift in proportion in the oral community in response to a suspension of OHP. In the search for microbial candidates for perio-probiotics, the taxa observed here to significantly decrease in the absence of OHP may prove to provide clinical benefits if supplemented to those at risk of progressing to periodontal disease. On the other hand, taxa observed to increase may be useful diagnostic indicators and therapeutic targets. Our reference-mapped variants contribute new candidates for these purposes.

Recently,Marotz et al. proposed the log-ratio of *Treponema* to *Corynebacterium* in saliva as a robust diagnostic indicator for periodontal disease. In this work, we found that *Corynebacterium* species observed mixed dynamics after the suspension of OHP. A rare significant result from saliva, *Corynebacterium matruchotii* increased in this niche (Figure S5) while *Corynebacterium durum* significantly decreased in the subgingival and supragingival plaque. The significant results from *Treponema* were in the sub- and supragingival plaques, where *Treponema vincentii*, *medium*, and *maltophilum* increased significantly in subgingival and *Treponema socranskii* and *vincentii* in supragingival plaque (Figure S1).

## Methods

### Participant demographics and sample collection

For this longitudinal study, 15 subjects (seven males and eight females aged 19 to 29 years) were recruited through the University of Toronto, Faculty of Dentistry as per guidelines of the Research Ethics Board (REB 30044). All had given informed consent to participate and followed the complete trial design. Oral health status of participants was determined by a dentist who performed a full mouth clinical examination that included the inspection of teeth, oral mucosa and periodontal status. All subjects were systemically healthy, with no active or previous history of periodontal disease, chronic aphthous ulcers or tonsillitis, and no more than 4 cavitated lesions. The absence of periodontitis was clinically assessed on all patients through analysis of tissue inflammation, bleeding on probing scores, and absence of periodontal destruction including clinical attachment loss. None were pregnant, past or current smokers, or users of non-steroidal anti-inflammatory or anti-microbial drugs, mouthwashes, or vitamin supplements within the previous three months.

Each individual had seven study visits (Figure 6). All plaque samples were collected at the beginning of each visit, before any interventions or assessments were conducted. Only one experienced and trained clinician was employed to collect all samples, thereby maximizing consistency and minimizing inter-individual discrepancy. The pre-trial preparation took place 7 days prior (day −7) to baseline (day 0), where an oral exam, scaling, and cleaning were performed and oral hygiene instructions were given for the next 7 days. At day 0, subjects were asked to abstain from all oral hygiene practices, including chewing gum, for a period of 21 days. During the induction phase (day 0, day 7, day 14, and day 21) clinical assessment involving full mouth pocket depth measurements and bleeding on probing (BOP) were performed on all teeth except the third molars. The clinical examinations were done using a University of North Carolina Probe. Gingival index (GI) (Löe and Silness, 1963) and plaque index (PI) (Löe and Silness, 1963; Löe, 1967) were recorded for the 6 surfaces of each tooth (disto-buccal, middle, mesio-buccal, mesio-lingual/mesio-palatal, middle, disto-lingual/disto-buccal). Oral rinse and blood samples were also collected to be used for another study investigating the relationship between inflammatory host response and gingivitis. At the end of day 21, subjects received professional prophylaxis, and were instructed to resume normal hygiene practices. They also received toothbrushes to facilitate the resolution of gingival inflammation. For another 2 weeks – at day 28 and day 35 – subjects underwent final clinical assessments and sample collection to confirm restoration of gingival health.

**Figure 6.**
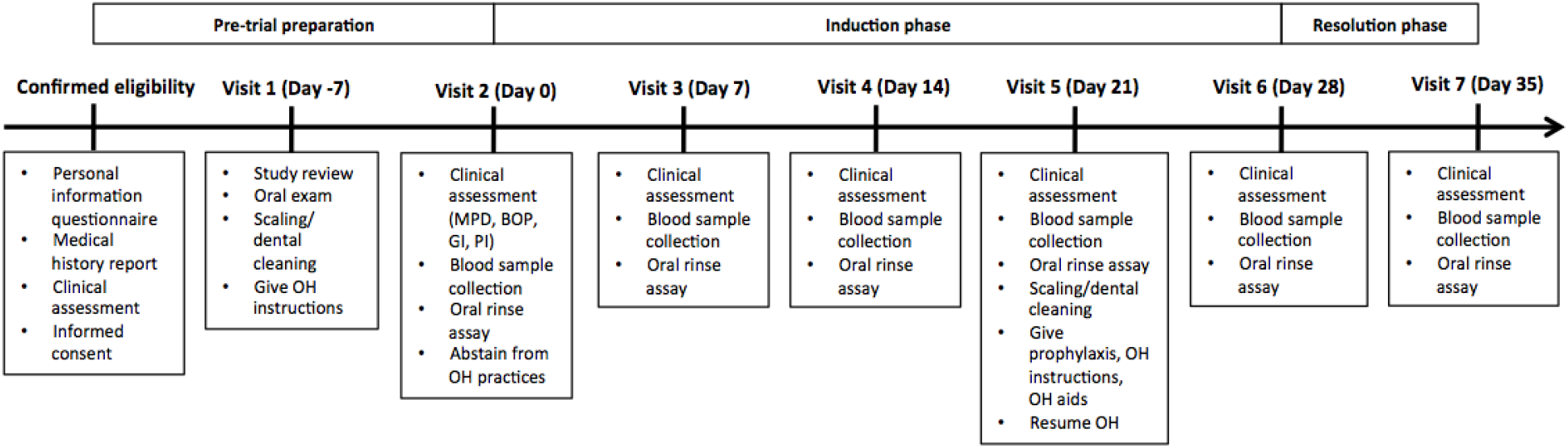
Study timeline and measurements at each visit. OH: oral hygiene; MPD: mean pocket depth; BOP: bleeding on probing; GI: gingival index; PI: plaque index

Collection of all oral samples was performed by the same clinician. Subgingival, supragingival, and tongue samples were collected by using separate dental curettes. Subgingival and supragingival plaques were obtained from Ramford’s 6 teeth (Griffen et al., 2011; Könönen and Wade, 2015; Zilm et al., 2007; Coit et al.,2016; Kreth et al., 2008). Plaque from tongue was collected by carrying out midline scrapings for a maximum of 4 times along the dorsum of the tongue using sterile inoculation loop. Stimulated salivary samples were collected in conical tubes and submerged on ice. Participants were asked to chew on a piece of plastic parafilm for 30s and subsequently swallowed the first saliva. Afterwards, participants expectorated once every 30s and a total of 4-5ml of saliva was collected each time into the conical tube. Salivary flow rate was calculated by dividing the total volume of saliva accumulated by the time required for collection. Saliva was aliquoted into sterile 1.5ml microcentrifuge tubes (Eppendorf; Axygen, CA, USA) for storage at −80°C until further processing.

### DNA extraction, sequencing, and quantification

DNA was extracted from 500μl of plaque using the DNeasy PowerSoil Kit (QIAGEN) according to the manufacturer’s instructions, with the exception of several modifications. The plaque sample was homogenized by 15 passages through a 3ml syringe with an 18G x 1.5” needle prior to extraction. The PowerBead Tubes were vortexed using Thermo Savant FastPrep FP120 Cell Homogenizer at 5 m/s for 45s. Samples were divided into two tubes prior to adding Solution C3 and subsequently mixed into a single spin filter. Final elution of DNA was completed with 150μl of Solution C6. DNA concentrations were determined by UV spectrophotometry (Ultrospec 3000, Pharmacia Biotech) at a wavelength of 260nm.

Amplification and sequencing of the 16S rRNA gene was performed as described in Comeau *et al*. (2017) (Comeau et al., 2017) and Walters *et al*. (2016) (Walters et al., 2016). Briefly, a one-step PCR approach is used to attach dual indexes, adaptors, and primers, followed by PCR amplification in duplicate, pooling, and clean up and normalization with the Invitrogen SequalPrep kit. Primers used were 515FB (5’-GTGYCAGCMGCCGCGGTAA-3’) and 926R (5’-CCGYCAATTYMTTTRAGTTT-3’) which span the V4-V5 hypervariable regions. Isolated and amplified DNA was then sequenced on an Illumina MiSeq to produce 300bp+300bp paired-end reads.

### Computational analysis

Sequences were processed using the QIIME2 software library (Bolyen et al., 2019). Sequence filtering, denoising, merging, and amplicon sequence variant (ASV) clustering was completed using DADA2 (version 1.22.0) via the q2-dada2 plugin (Callahan et al., 2016). A number of nucleotides the length of the forward and reverse primers were trimmed from their respective reads before assembly. Alpha diversity analyses were performed with the q2-diversity plugin. Phylogenetic trees were generated using MAFFT (version 7.490) via the q2-phylogeny plugin (Katoh and Standley, 2013). Taxonomic classifications were assigned with the q2-feature-classifier using the full-length, 99% pre-clustered SILVA 138 pretrained Naïve Bayes classifier artifact (Pruesse et al.,2007).

To account for the compositional nature of the data, the ASV table and collapsed taxonomic tables were transformed using a centered log ratio (CLR) transform before analyzing for trends using the compositions R library (version 2.0-4) (Van den Boogaart and Tolosana-Delgado, 2008). This allowed for trends during the induction phase to be modeled using a generalized linear model (GLM) via the lme4 toolkit (version 1.1-29) (Bates et al., 2014). The model used is described by the lme4 formula `CLR_abundance ~ time + (1|subject)` describing a repeated measures experiment, which fits a slope to the transformed relative abundances as a function of time. Significant slopes were assessed with the anova.lmerModLmerTest function from the lmerTest package (version 3.1-3) (Kuznetsova et al., 2017), which uses Satterthwaite’s method to estimate degrees of freedom and F-statistics. Multiple hypothesis test correction was completed using false discovery rate (FDR) correction. Phylogenetic trees were visualized using ggtree (version 3.2.1) (Yu, 2020).

## Conclusions

The EGM is a valuable experimental system to correlate gingivitis onset and its resolution with various physiological and ecological parameters in the human mouth. Although gingivitis includes varying levels of gum inflammation, all forms are characterized by swollen, dark red and tender gums that bleed easily during brushing and flossing. Taking advantage of the fact that 100% of participants who completely restrain from daily oral hygiene for three weeks will develop gingivitis, here we utilized the EGM with 16S rRNA gene sequencing to document comprehensive snapshots of oral bacterial succession in four distinct oral habitats. Through phylogenetic placement of ASVs alongside eHOMD reference sequences, we reveal a high-resolution view of relative shifts, wherein oral taxa whose abundance were significantly increased (i.e., in the case of putative and known pathogens) or decreased (i.e., commensals or health-associated microbes) by the end of disease-induction (on Day 21) were identified.

We demonstrated that microbial abundance shifts were largely reversed after daily oral hygienic practices were re-introduced in the final two weeks of the study. This study provides a useful phylogenetic reference with ASVs and species from this study aligned to and highlighted in the eHOMD reference set. Researchers can make use of the phylogenetic atlas presented here to identify taxa of interest, and if sufficient resolution exists within the V4-V5 16S rRNA gene fragment for profiling their microbial taxa or complexes of interest when designing future studies. While it is yet to be investigated if some subset of these bacterial markers can be monitored to predict the onset of periodontal disease if gingivitis is left untreated, such studies could create a path for early detection and intervention strategies against periodontitis before irreversible bone damage is caused.

## Supplementary Information

## A. Appendix I

### Interpreting compositional abundance shifts

The underlying abundance data in this study are gene counts, which are positive integer values that are often transformed prior to analysis and interpretation. Gene/taxon abundance data, when presented as raw counts or expressed as proportions of the sample, are compositional in nature (Gloor et al., 2017). There are significant and ongoing discussions about the impact this property has on the use and interpretation of community gene abundance data and the appropriate application of statistical methods (Quinn et al., 2019). The primary limitation to highlight for this work is the difficulty in interpreting the direction of change in proportional or percent abundance data. For example, as a percentage of the subgingival community, the genus *Streptococcus* decreased from 17.89% to 3.91%, while a simultaneous increase is observed in *Prevotella* which increased from 21.63% to 40.12%, on average. Without additional validation such as quantitative PCR analysis, there is an unresolvable ambiguity about whether the near-doubling of *Prevotella* is, in fact, a near-doubling of the number of *Prevotella* cells in the environment or whether a decline in other community members (such as *Streptococcus*) is leading to the *appearance* of an increase in *Prevotella* proportional abundances. In reality, it is likely some combination of those two possibilities and our observed abundances are the result of that push and pull.

In this study, two types of abundance data are routinely presented: proportional/percent abundances of gene counts and CLR-transformed gene counts. The former are generally more intuitive values that represent the proportion of a whole, while the latter transforms all counts as a log-ratio using the geometric mean of the sample as the denominator. While the CLR-transform benchmarks all taxon abundances relative to the average taxon in that sample and does not place the transformed values on a simplex as proportional/percentage values are, the interpretation of CLR-transformed values is less intuitive. The CLR-transformed abundances can be interpreted as the fold-difference in abundance between the scrutinized taxon and the average taxonomic unit at that level. A CLR-transformed value of 0 implies that the taxon has the same abundance as the average taxonomic unit (after the addition of pseudocounts). Positive values indicate a larger fold difference compared to the community average and negative values indicate taxonomic counts that are lower than the average taxon in the sample.

For these reasons stated above, statements about the absolute abundance of the organisms described in this study cannot be made with this data and to do so would require additional laboratory analysis. However, the pre-EGM and recovery phase samples represent important benchmarks that enable comparisons of the overall importance of each taxonomic unit relative to the average taxon in the community. When these benchmarks are compared to the weekly time points collected during the induction phase, the temporal trajectory of each taxon after suspension of OHP, in terms of its relative weight in the community, can be stated.

### Descriptions of bacterial dynamics across phyla

#### Spirochaetota

Supragingival and subgingival plaque samples showed significant increases in CLR-transformed abundances of reads classified to the phylum Spirochatota over the gingivitis induction period (Figure 4). The vast majority of reads classified to this phylum were further classified to the genus *Treponema*, which showed the same significantly increasing trends in supra- and subgingival plaque samples (Figure 5). At the species level, six species demonstrated significant trends over gingivitis induction, with subgingival levels of *T. medium*, *T. socranskii*, and *T. maltophilum* almost doubling in mean relative abundance. The lower abundance species *T. vincentii* increased nearly 6-fold from 0.03% to 0.19% mean relative abundance in the subgingival plaque, and doubled its mean abundance in supragingival plaque from 0.03% to 0.06%. The two remaining significant species, *T*. genomosp. and *T. refringens*, were ultra-low abundance (<0.01%) but nonetheless saw a significant decrease in the supragingival and subgingival plaque, respectively (Figure S1, Table S1). At the ASV level, three variants increased over the induction period: one *T. vincentii* and one *T. medium* ASV increased in subgingival plaque, and one *T. socranskii* ASV increased in the supragingival plaque (Figure S1).

#### Pseudomonadota

The phylum Pseudomonadota decreased sharply and significantly in the tooth plaque samples after the first week of induction (Figure 4), driven by the most abundant genus *Neisseria* (Figure 5), which accounted for 9.58% and 12.8% on day 0 in the subgingival and supragingival plaque, respectively, and fell significantly to 2.23% and 4.96% by day 21. Interestingly, *Neisseria* increases in abundance for the first two weeks after suspension of OHP before rapidly falling for an overall decrease in community representation. An ASV classifying to *Kingella*,a close neighbour of *Neisseria*, decreased significantly in supragingival plaque samples over the induction period (Figure S4).

In the subgingival plaque, *Haemophilus* showed a 30-fold decrease over the gingivitis induction period, from 3.41% to 0.11% mean read abundance and a 5-fold decrease from 3.55% to 0.63% in the supragingival plaque. Three ASVs that classified to *Haemophilus* and all placed closely on the reference tree to *H. parainfluenzae* had significantly decreasing trends in the subgingival plaque, with one of the ASVs also decreasing significantly in the supragingival plaque (Figure S2). Similarly, sequences from the genus *Lautropia* decreased 3-fold in both supra- and subgingival plaque, though the result was only statistically significant for the supragingival plaque (Figures 5, S3).

In contrast with the generally decreasing trends in this phylum, the genus *Aggregatibacter* demonstrated an increasing trend in both tooth plaques and on the tongue. This genus was the sole taxonomic unit (whether phylum, genus, species, or ASV) that significantly trended in either direction on the tongue. In both tooth plaque environments, *Aggregatibacter* almost doubled their relative abundance from about 0.8% to 1.5%, but increased almost 7-fold on the tongue from 0.014% to 0.093% (Figure S2).

Located between *Aggregatibacter* and *Neisseria* on the reference tree, three additional ASVs increased significantly over the induction phase. In aggregate, *Cardiobacterium valvarum* sequences increased significantly in both tooth plaque environments, while a specific *C. valvarum* ASV increased significantly only in the supragingival plaque samples. An ASV that classified as *Comamonadaceae* and placed on the reference tree near genus *Ottowia* increased in the supragingival plaque. The final significant ASV in this group classified to *Propionivibrio*, placed near genus *Rhodocyclus*, and increased in the subgingival plaque (Figure S3).

#### Actinomycetota

Abundances of Actinomycetota fell quickly and significantly after OHP were suspended (Figure 4, Table S1). This was driven by a significant 4.2 and 3.8-fold decrease in abundance of the genus *Actinomcyes* in the subgingival and supragingival plaque, respectively (Figure 5, Table S1). Three ASVs in this genus significantly decreased in tooth plaque samples, including two that placed on the reference tree near a group of sequences with low nucleotide variation in the sequenced V4-v5 region that originated from *A. oris*, *A. naeslundii*, *A. viscosus*, and *A. johnsonii*. The third ASV placed most closely to *A. gerencseriae* and decreased significantly in the subgingival plaque.

The second-most abundant genus in the phylum was *Rothia*, which decreased in both the subgingival and supragingival plaque, seeing a significant 20.8 and 8.8-fold decrease, respectively. In the subgingival plaque, *Rothia* mean abundance fell from 1.24% to 0.06%, with supragingival plaque mean abundance similarly dropping from 1.62% to 0.18%. Two ASVs that classified to *Rothia* significantly decreased over the induction phase. The first decreased significantly in the subgingival plaque and placed on the reference tree most closely to *R. aeria*. The second decreased significantly in both plaque types and placed most closely to *R. dentocariosa*.

A low-abundance uncultivated group F0332 in the family *Actinomycetaceae* decreased significantly in the supragingival plaque from 0.36% to 0.07%, with a concurrent 8-fold, but not statistically significant, decrease in the subgingival plaque. This was driven by an ASV that placed with sequences belonging to genus *Peptidiphaga* When sequence abundances were aggregated at the genus level, *Corynebacterium* abundance increased significantly only in the saliva, driven by sequences that classified down to *C. matruchotii*. More in line with other members of its phylum, *C. durum* decreased in abundance in both tooth plaque sets, led by one primary ASV (Figure S5, Table S1). (Figure S6, Table S1). One final very low abundance ASV that placed near members of the *Arachnia* genus increased significantly in the fourth week of induction (Figure S5).

#### Bacillota

The phylum Bacillota is a large and diverse group, so while it decreased significantly in abundance in the subgingival plaque after the first full week of the cessation of OHP (Figure 4), it also contains several lower abundance members that increased significantly in abundance over the same period. The phylum-level decrease following the first week was driven primarily by a reduction of the most abundant genus, *Streptococcus*, and to a lesser extent *Granulicatella*. On the other hand, *Selenomonas*, *Dialister*, *Johnsonella*, *Gemella*, *Parvimonas*, *Catonella*, *Centipeda*, and *Abiotrophia* are all low abundance genera that saw significant increases over gingivitis induction (Figure 3, Table S1). Bacillota also contains one of the most abundant genera that did not significantly increase or decrease in any site, *Veillonella*.

Over the induction period, *Granulicatella* saw its relative proportion in the community shrink by a factor of 4 in both the tooth plaque environments. Meanwhile, neighbouring genus *Abiotrophia*, increased its representation in saliva samples by 4-fold, and *Gemella* increased significantly in the supragingival plaque but began declining by week 4 of suspended OHP (Figure S7). One ASV drove the *Granulicatella* decreases, most closely placing on the reference tree near *G. adiacens*, while another ASV drove the *Abiotrophia* increase in saliva, placing close to an *A. defectiva* reference sequence (Figure S7).

Members of the genus *Streptococcus* saw a steady decrease after suspension of OHP in both tooth plaque environments, with a 4.6-fold decrease in subgingival plaque and 2.9-fold decrease in supragingival plaque by day 21. Three streptococcal ASVs decreased significantly in subgingival plaque, falling on the reference tree near *S. oralis/S. mitis, S. parasanguinis/S. australis*, and *S. salivarius/S. vestibularis/S. thermophilus*. A final ASV, high in abundance and placing near *S. sanguinis*, decreased significantly in both the subgingival and supragingival plaques (Figure S8).

The genus *Selenomonas* increased significantly in the tooth plaque, increasing from 1.9% of subgingival samples at day 0 to 3.1% at day 21. The closely related and lower abundance genus *Centipeda* also increased in the tooth plaque samples over the induction phase. Two of the significant *Selenomonas* ASVs increased significantly in saliva samples, placing near *S. infelix* and *S. artemidis*. At the species and ASV level, *S. sputigena* sequences increased significantly in the tooth plaque samples and accounted for nearly half of *Selenomonas* sequences in subgingival plaque at day 21 (Figure S10, Table S1).

Additional low abundance members of the *Bacillota* phylum include *Peptococcus* and *Dialister* (Figure S9), *Catonella, Johnsonella, Lachnoanaerobaculum*, and Clostridia UCG-014 (Figure S11), and *Parvimonas* (Figure S12), which were generally found to be increasing in the tooth plaque samples, particularly subgingival samples.

#### Patescibacteria & Campylobacterota

*Patescibacteria*, sometimes known as SR1, is a very low abundance group that showed significant increases in the tooth plaque samples, with the *Absconditabacteriales* group increasing from 0.008% of reads mapped to 0.031% by day 21 in subgingival samples. One ASV in this group significantly increased in supragingival samples, and mapped most closely to HMT 345 in the eHOMD reference database (Figure S14). A genus-level designation JGI 0000069-P22 increased significantly in both tooth plaque samples, a result that was significant only in supragingival plaque at the species level as Gracilibacteria bacterium, and for an ASV that placed near eHOMD HMT 872 (Figure S13).

The phylum *Campylobacterota* increased significantly in the tooth plaque samples. At the species level, *C. showae* sequences increased significantly in the supragingival plaque (Figure S13).

#### Fusobacterota

As a phylum, the *Fusobacterota* doubled their mean reads mapped from day 0 to day 21 in both tooth plaque environments. This was driven by members of the genus *Fusobacterium*, which doubled from 8.6% to 17.4% in the subgingival plaque, and the genus *Leptotrichia*, which doubled from 2.0%to 4.1% in subgingival plaque.

*Fusobacterium* sequences included 7 ASVs that increased in the tooth plaque, and one that increased in the tooth plaque and saliva samples. The ASV that increased significantly in all three environments placed most closely to *F. nucleatum* subsp. *polymorphum* (HMT 203). Tooth plaque ASVs placed near *F. nucleatum* subsp. *vincentii* (HMT 200, 205), *F. hwasookii* (HMT 370, 953), and *F. nucleatum* subsp. *animalis* (HMT 420) (Figure S15).

At the species level, *Leptotrichia* saw significant increases in reads mapping to *L. buccalis, L. shahii*, and a significant decrease in supragingival plaque in low abundance species-level designation Leptotrichia-like sp. (Figure S16). The *L. buccalis* increase was driven by an ASV that mapped near HMT 563, and similarly the *L. shahii* with HMT 214. Additional *Leptotrichia* ASVs that placed near HMT 219 and HMT 417 increased significantly in the subgingival plaque (Figure S16).

#### Bacteroidota

The *Bacteroidota* are another large phylum with a diverse set of dynamics exhibited after the suspension of OHP. As a phylum, its members nearly double their community proportions by day 21, accounting for a mean of 40% of subgingival samples at this time point. More than half of this abundance belongs to the genus *Prevotella* which increased significantly in both tooth plaque environments. The genera *Porphyromonas*, *Alloprevotella*, *Tannerella*, and genus-level designation F0058 also increased significantly in the tooth plaque samples.

The genus *Prevotella* accounted for an average of 8.9% of reads mapped at day 0 and increased to 21.0% by day 21 in the subgingival plaque with a similar increase observed in the supragingival plaque. At the species level, a wide number of named *Prevotella* species increased in one or both of the tooth plaque environments and saliva samples:

*P. shahii*, *P. nigrescens*, *P. loescheii*, *P. saccharolytica*, *P. micans*, *P. maculosa*, *P. oulorum*, and *P. marshii*, while *P. histicola* sequences saw their salivary proportion decrease significantly from 1.9% at day 0 to 0.7% at day 21.

The ASV-level results reveal the same diversity seen in the species-level aggregations. In total 9 ASVs classifying to *Prevotella* increased significantly in the tooth plaque and/or saliva. Three of these ASVs placed near the significant species-level designations *P. micans*, *P. nigrescens*, *P. shahii*, and *P. saccharolytica*. Of the remaining 5 ASVs, two placed near *P. melaninogenica*, and eHOMD reference sequences HMT 475 and HMT 301. The two *P. melaninogenica* ASVs include one that decreases significantly over time and another that increases, in a rare example of opposing dynamics in such closely related units (Figure S19).

The second-most abundant genus in the *Bacteroidota* was *Capnocytophaga*, which saw non-significant increases in abundance across all sample types. The only named species with a significant increase was *C. granulosa*, which increased significantly in supragingival plaque. Counter to the dynamics in the genus, *C. gingivalis* and *C. leadbetteri* sequences decreased significantly in subgingival plaque, though this followed a period of initial increase after the suspension of OHP. Nearby genus *Bergeyella* had species-level designation “uncultured *Bergeyella”* increase in both tooth plaque types and saliva samples (Figure S17).

Members of the genus *Porphyromonas* doubled their proportion of the tooth plaque community, on average, over the induction phase. A similar increase was seen in the saliva but was not statistically significant after p-value correction. At the species level, *P. catoniae* significantly increased over two-fold in the subgingival plaque and over 5-fold in supragingival plaque. One ASV matched the dynamics of the species-level aggregate for *P. catoniae* and increased significant in both tooth plaques and saliva. A second significant *Porphyromonas* ASV increased from low abundance after suspension of OHP in supragingival plaque and saliva, placing near eHOMD HMT 275, HMT 277, and HMT 284 (Figure S18).

*Alloprevotella* increase their proportion of the community nearly 5-fold in the subgingival plaque, from 0.8% to 4.0%, with a significant two-fold increase in supragingival plaque also observed. The species *A. tannerae* increased its proportion over 8-fold between day 0 and day 21 in the subgingival plaque. Two ASVs placed near *A. tannerae* (HMT 466) sequences and increased significantly in the subgingival plaque (Figure S18).

As a genus, *Tannerella* sequences increased roughly 3-fold in both tooth plaque environments. *T. forsythia* proportions decreased significantly in the supragingival plaque samples, from a mean of 0.05% to zero detection across all subject samples. Three *Tannerella* ASVs increased significantly in both tooth plaque environments, placing closer to HMT 286, HMT 916, and HMT 808 than the *T. forsythia* reference HMT 613 (Figure S18).

The genus-level designation F0058, belonging to the family *Paludibacteraceae*, increased significantly in both tooth plaque samples. While no one ASV in this group met the significance threshold, representative sequences from this taxonomic group placed near Bacteroidales G-2 bacterium (HMT 274). Unlike its phylogenetic neighbours, the low-abundance genus *Bacteroides* decreased significantly in both tooth plaque environments over the induction period (Figure S18).

---

**Figure S1.**
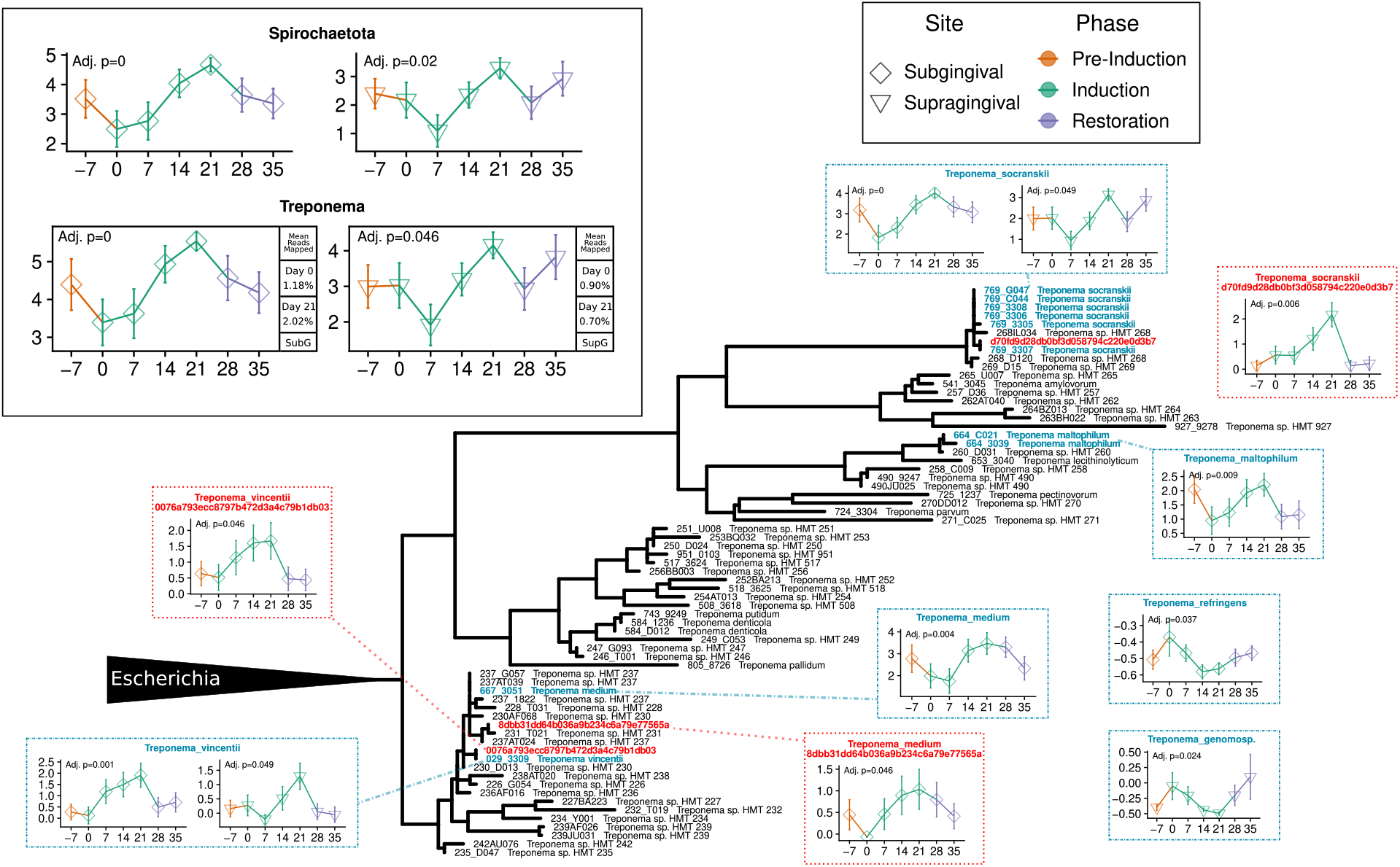
Treponema. Sequence abundances aggregated at the phylum and genus level (solid black outline), species level (blue dashed outline) and ASV level (red dotted outline). Y-axis is CLR-transformed reads and x-axis is days. Phylogenetic tree situates differentially abundant named species (blue) and ASVs (red) alongside the eHOMD reference sequences (black). p-values are the result of testing the null hypothesis that the slope is zero over the induction phase, with only significant results shown.

**Figure S2.**
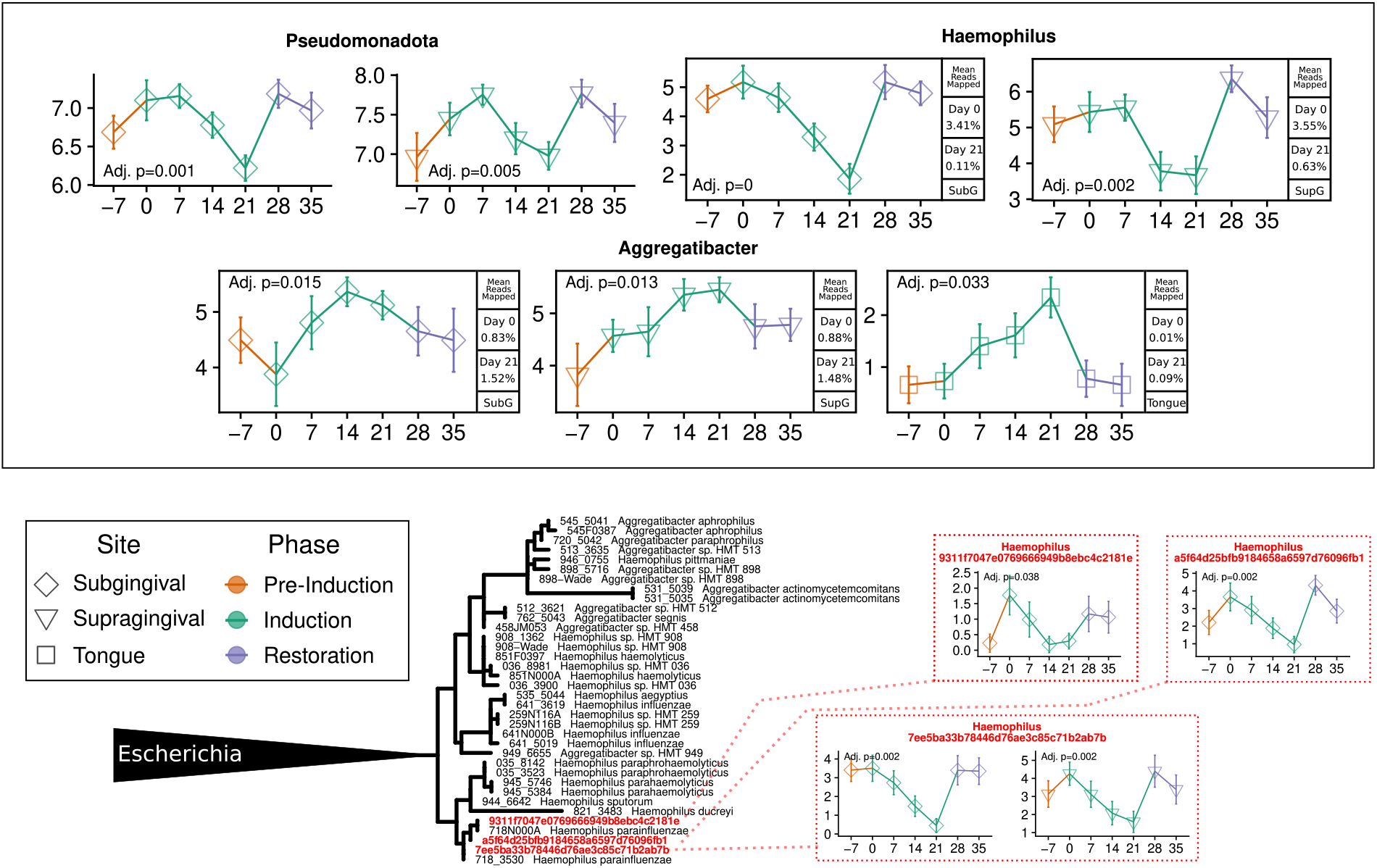
Haemophilus, Aggregatibacter. Sequence abundances aggregated at the phylum and genus level (solid black outline), species level (blue dashed outline) and ASV level (red dotted outline). Y-axis is CLR-transformed reads and x-axis is days. Phylogenetic tree situates differentially abundant named species (blue) and ASVs (red) alongside the eHOMD reference sequences (black). p-values are the result of testing the null hypothesis that the slope is zero over the induction phase, with only significant results shown.

**Figure S3.**
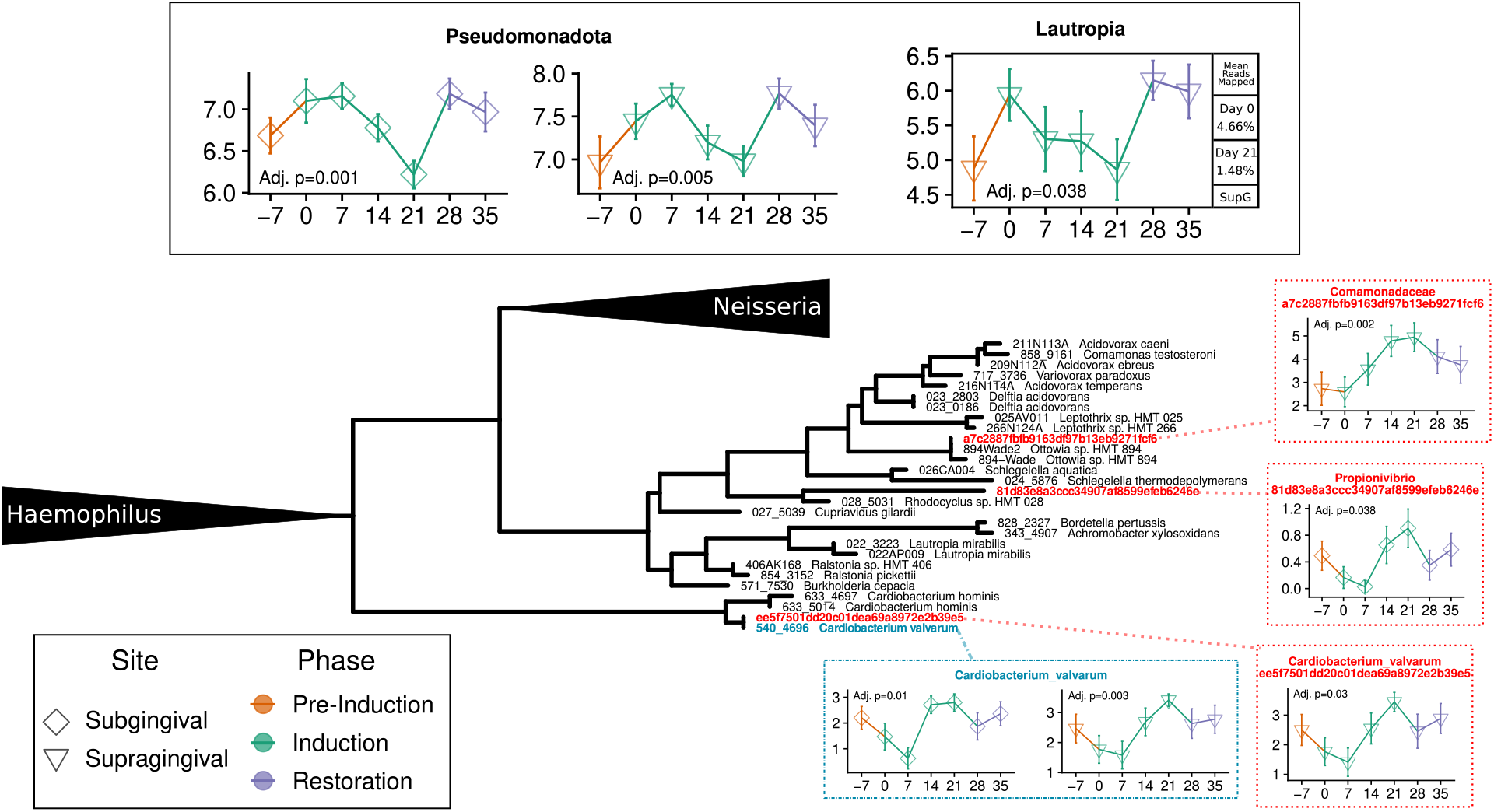
Lautropia, Cardiobacterium. Sequence abundances aggregated at the phylum and genus level (solid black outline), species level (blue dashed outline) and ASV level (red dotted outline). Y-axis is CLR-transformed reads and x-axis is days. Phylogenetic tree situates differentially abundant named species (blue) and ASVs (red) alongside the eHOMD reference sequences (black). p-values are the result of testing the null hypothesis that the slope is zero over the induction phase, with only significant results shown.

**Figure S4.**
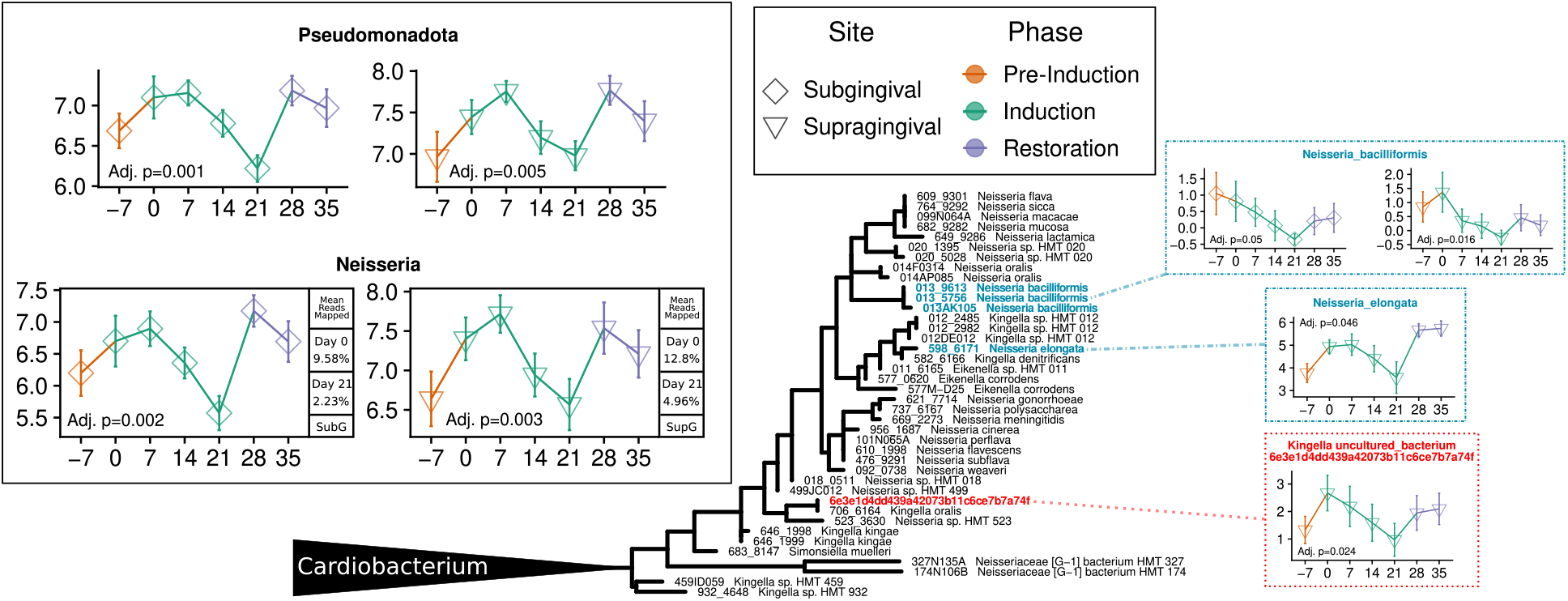
Neisseria, Kingella. Sequence abundances aggregated at the phylum and genus level (solid black outline), species level (blue dashed outline) and ASV level (red dotted outline). Y-axis is CLR-transformed reads and x-axis is days. Phylogenetic tree situates differentially abundant named species (blue) and ASVs (red) alongside the eHOMD reference sequences (black). p-values are the result of testing the null hypothesis that the slope is zero over the induction phase, with only significant results shown.

**Figure S5.**
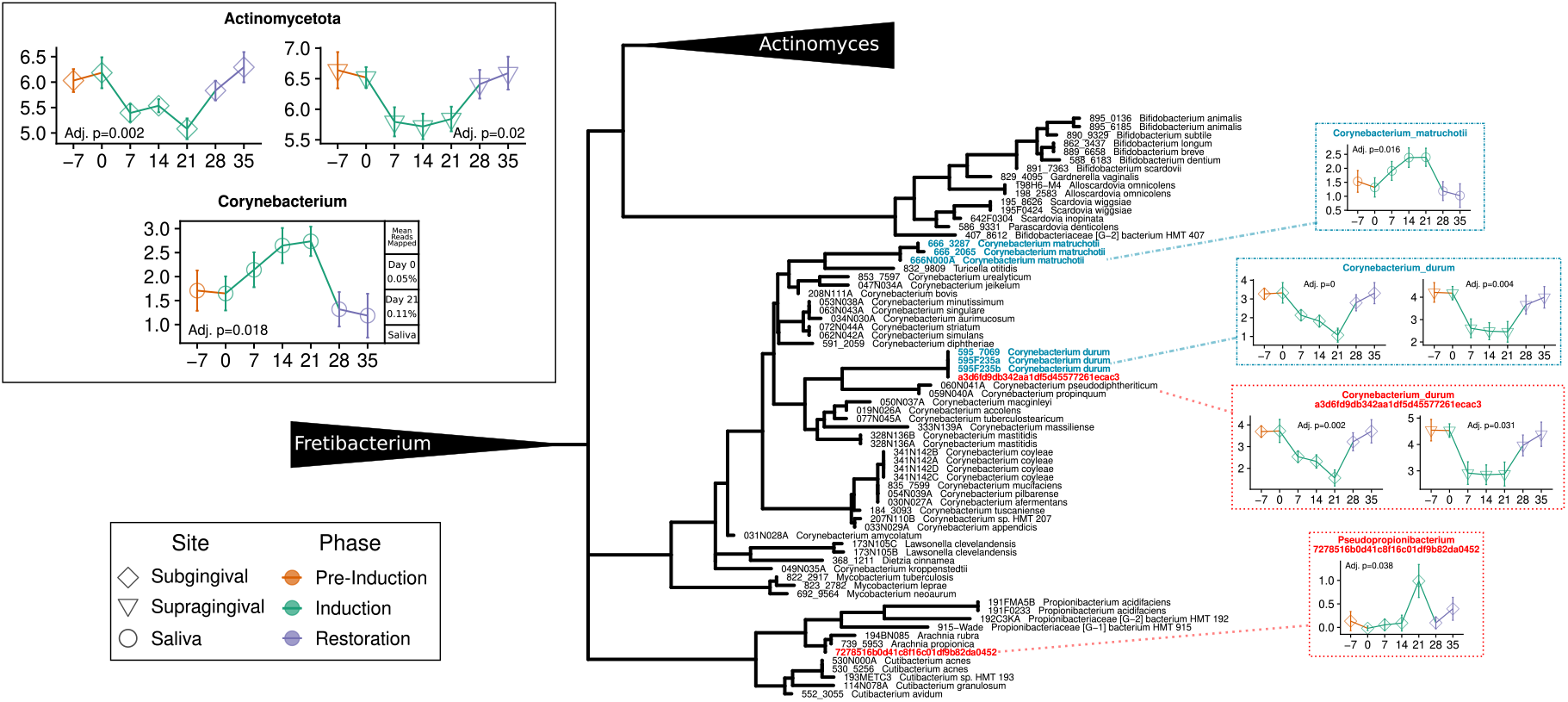
Corynebacterium. Sequence abundances aggregated at the phylum and genus level (solid black outline), species level (blue dashed outline) and ASV level (red dotted outline). Y-axis is CLR-transformed reads and x-axis is days. Phylogenetic tree situates differentially abundant named species (blue) and ASVs (red) alongside the eHOMD reference sequences (black). p-values are the result of testing the null hypothesis that the slope is zero over the induction phase, with only significant results shown.

**Figure S6.**
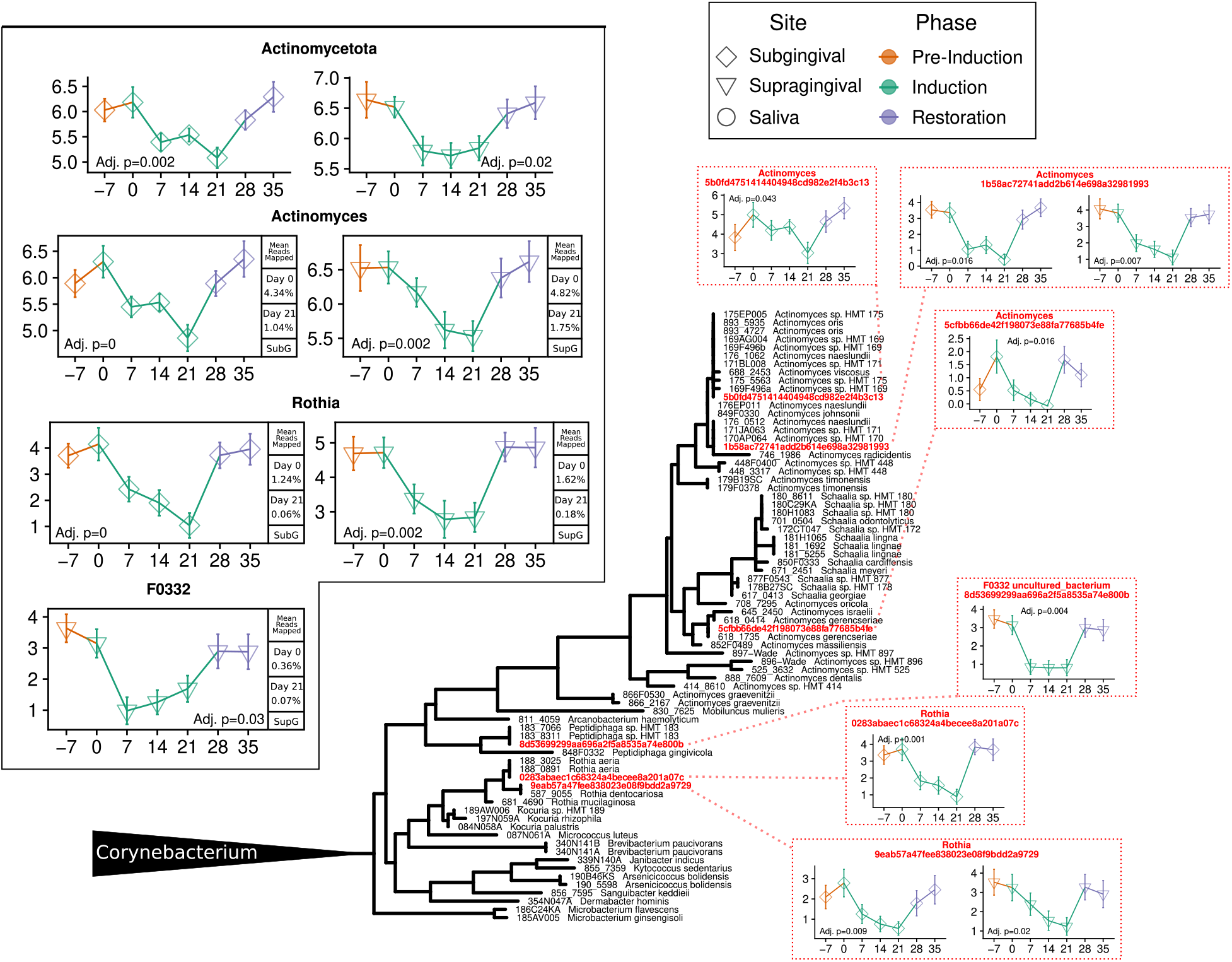
Actinomyces, Rothia. Sequence abundances aggregated at the phylum and genus level (solid black outline), species level (blue dashed outline) and ASV level (red dotted outline). Y-axis is CLR-transformed reads and x-axis is days. Phylogenetic tree situates differentially abundant named species (blue) and ASVs (red) alongside the eHOMD reference sequences (black). p-values are the result of testing the null hypothesis that the slope is zero over the induction phase, with only significant results shown.

**Figure S7.**
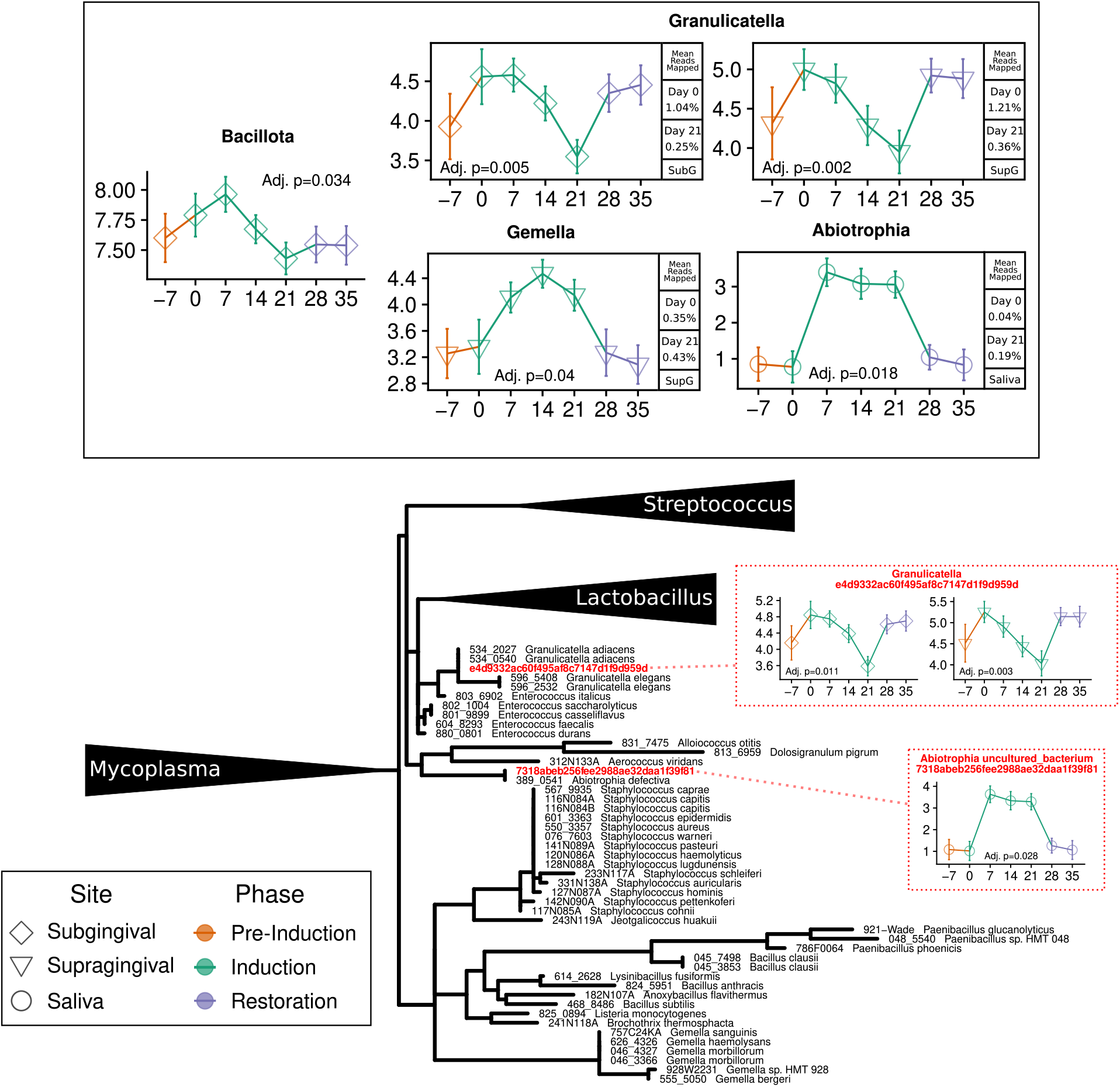
Granulicatella, Abiotrophia, Gemella. Sequence abundances aggregated at the phylum and genus level (solid black outline), species level (blue dashed outline) and ASV level (red dotted outline). Y-axis is CLR-transformed reads and x-axis is days. Phylogenetic tree situates differentially abundant named species (blue) and ASVs (red) alongside the eHOMD reference sequences (black). p-values are the result of testing the null hypothesis that the slope is zero over the induction phase, with only significant results shown.

**Figure S8.**
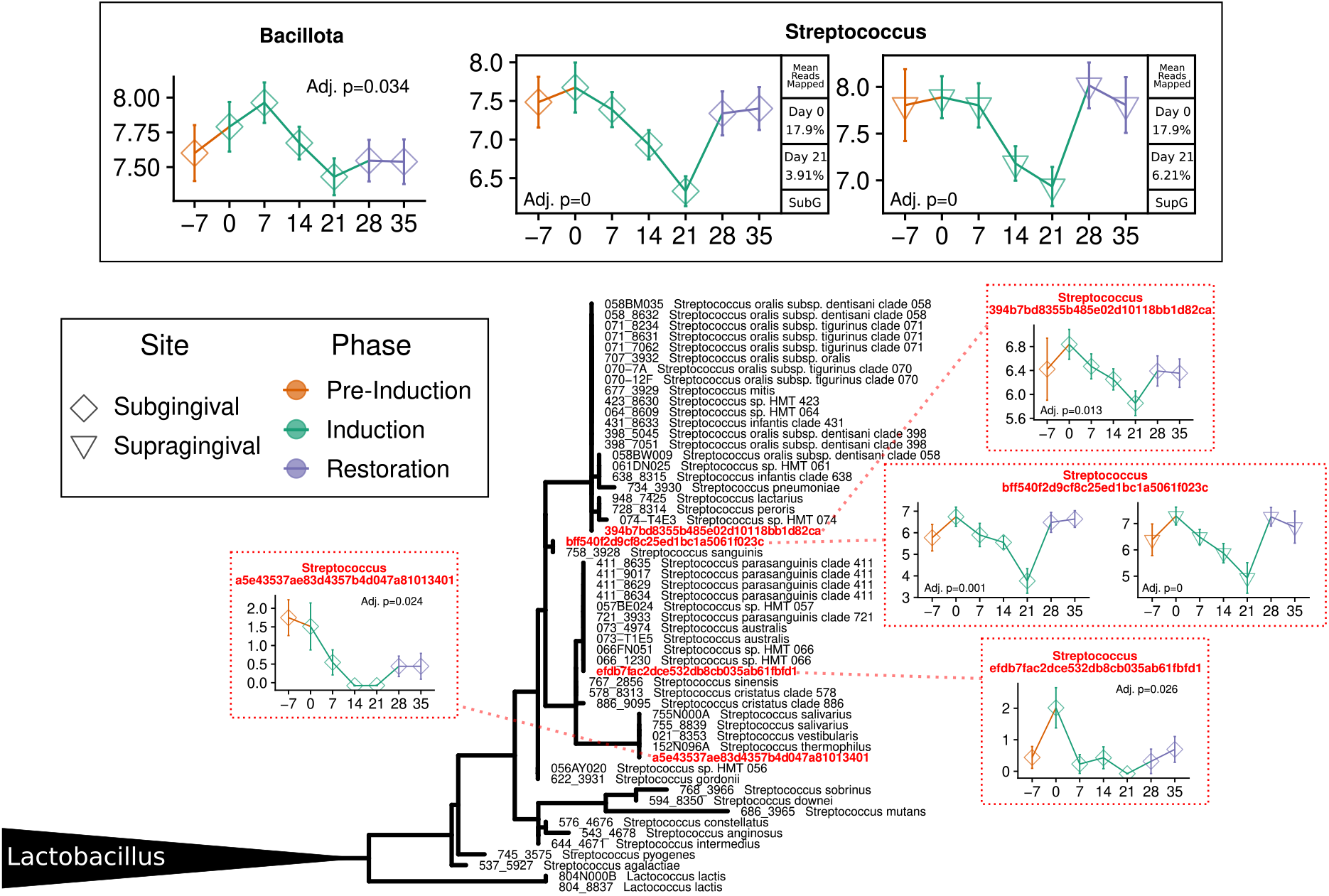
Streptococcus. Sequence abundances aggregated at the phylum and genus level (solid black outline), species level (blue dashed outline) and ASV level (red dotted outline). Y-axis is CLR-transformed reads and x-axis is days. Phylogenetic tree situates differentially abundant named species (blue) and ASVs (red) alongside the eHOMD reference sequences (black). p-values are the result of testing the null hypothesis that the slope is zero over the induction phase, with only significant results shown.

**Figure S9.**
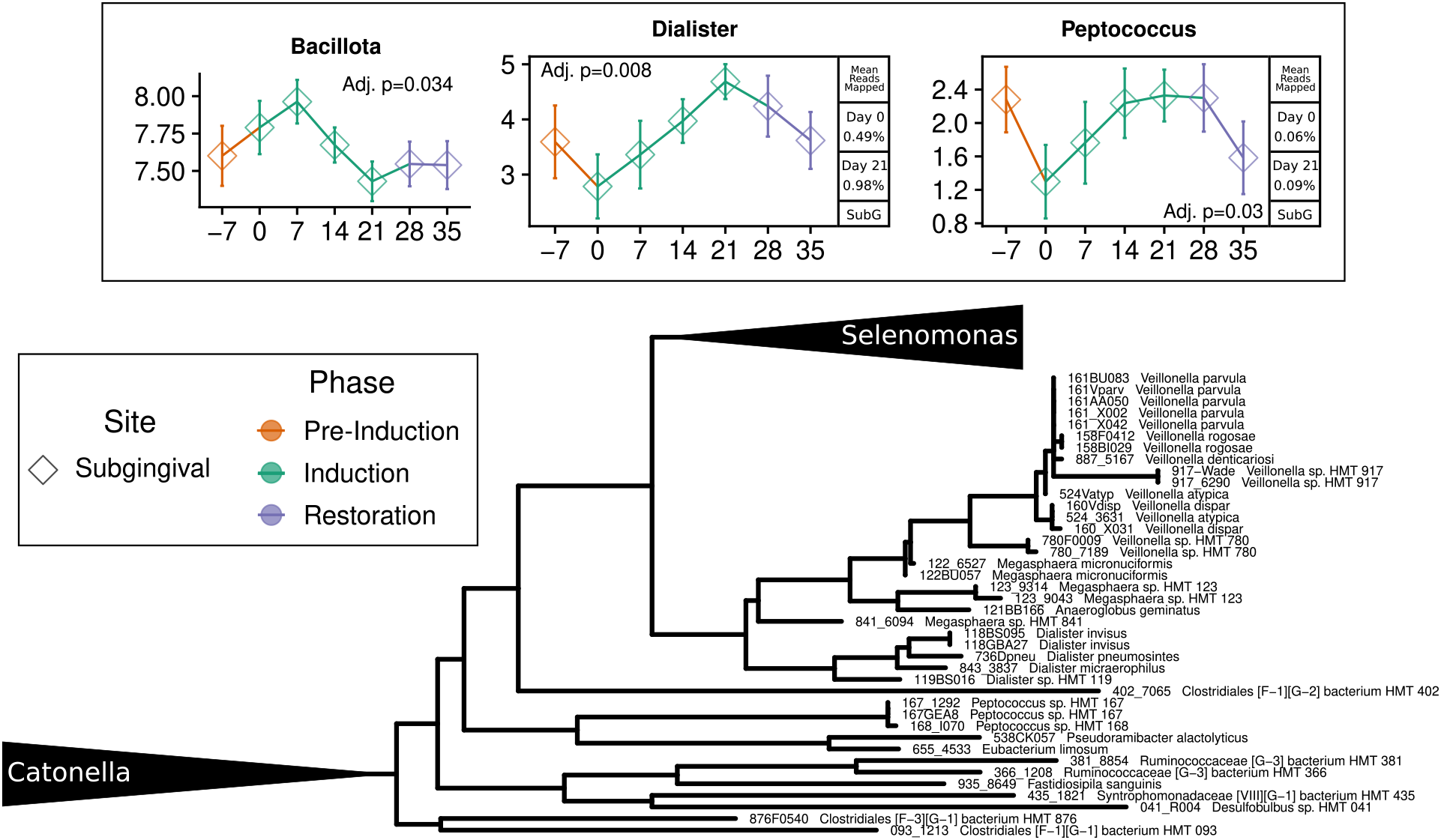
Veillonella, Dialister, Peptococcus. Sequence abundances aggregated at the phylum and genus level (solid black outline), species level (blue dashed outline) and ASV level (red dotted outline). Y-axis is CLR-transformed reads and x-axis is days. Phylogenetic tree situates differentially abundant named species (blue) and ASVs (red) alongside the eHOMD reference sequences (black). p-values are the result of testing the null hypothesis that the slope is zero over the induction phase, with only significant results shown.

**Figure S10.**
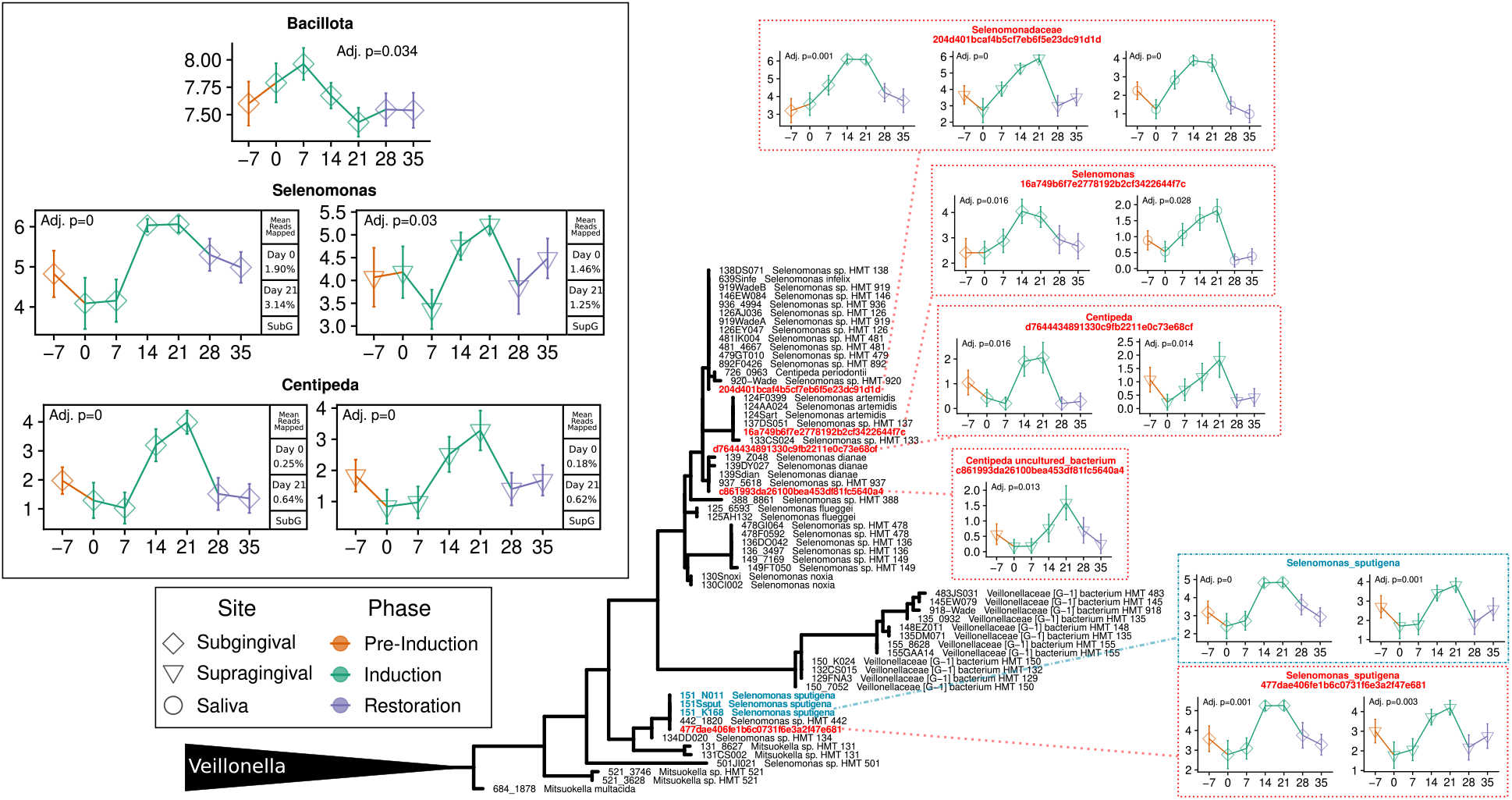
Selenomonas, Centipeda. Sequence abundances aggregated at the phylum and genus level (solid black outline), species level (blue dashed outline) and ASV level (red dotted outline). Y-axis is CLR-transformed reads and x-axis is days. Phylogenetic tree situates differentially abundant named species (blue) and ASVs (red) alongside the eHOMD reference sequences (black). p-values are the result of testing the null hypothesis that the slope is zero over the induction phase, with only significant results shown.

**Figure S11.**
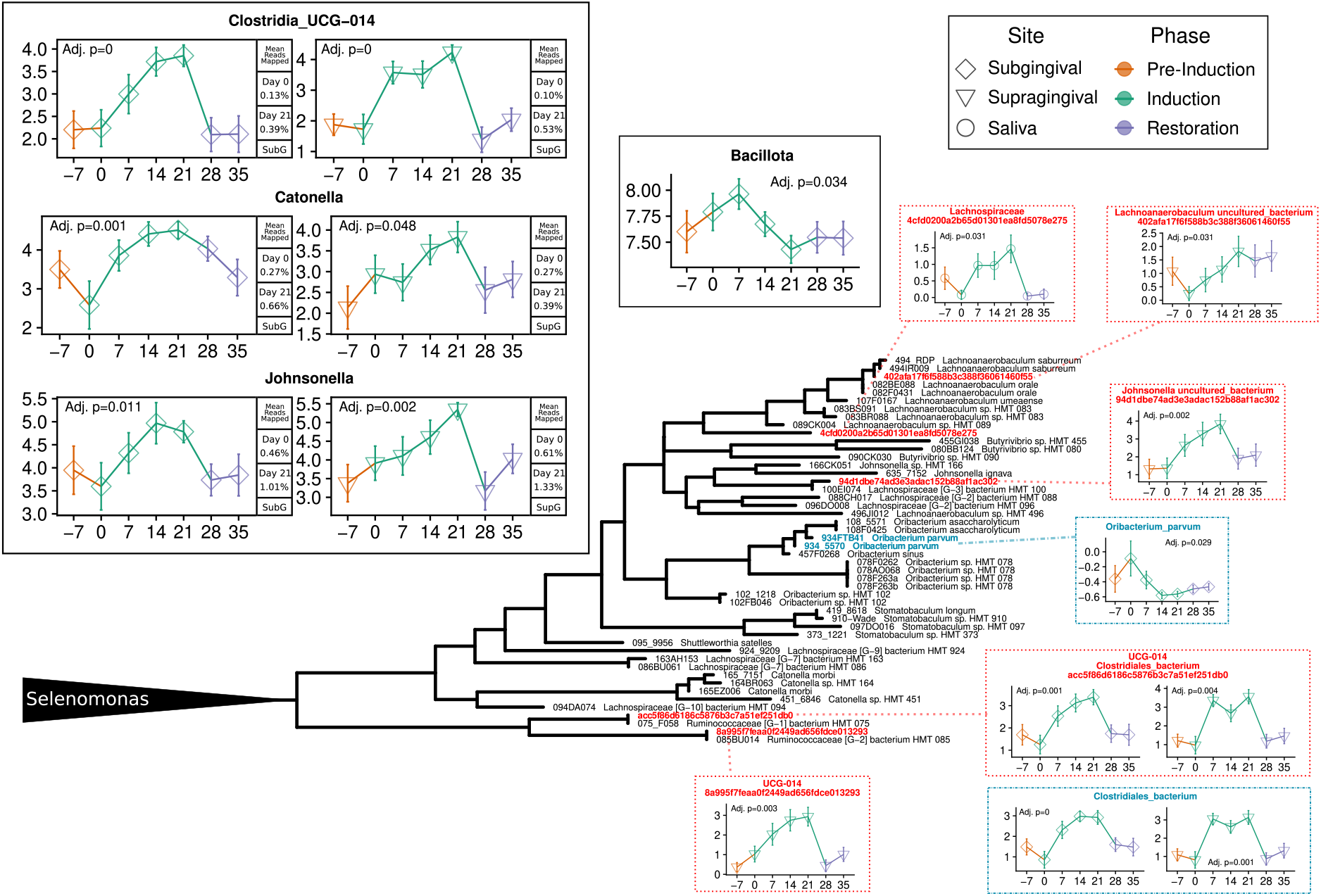
Catonella, Johnsonella, Oribacterium. Sequence abundances aggregated at the phylum and genus level (solid black outline), species level (blue dashed outline) and ASV level (red dotted outline). Y-axis is CLR-transformed reads and x-axis is days. Phylogenetic tree situates differentially abundant named species (blue) and ASVs (red) alongside the eHOMD reference sequences (black). p-values are the result of testing the null hypothesis that the slope is zero over the induction phase, with only significant results shown.

**Figure S12.**
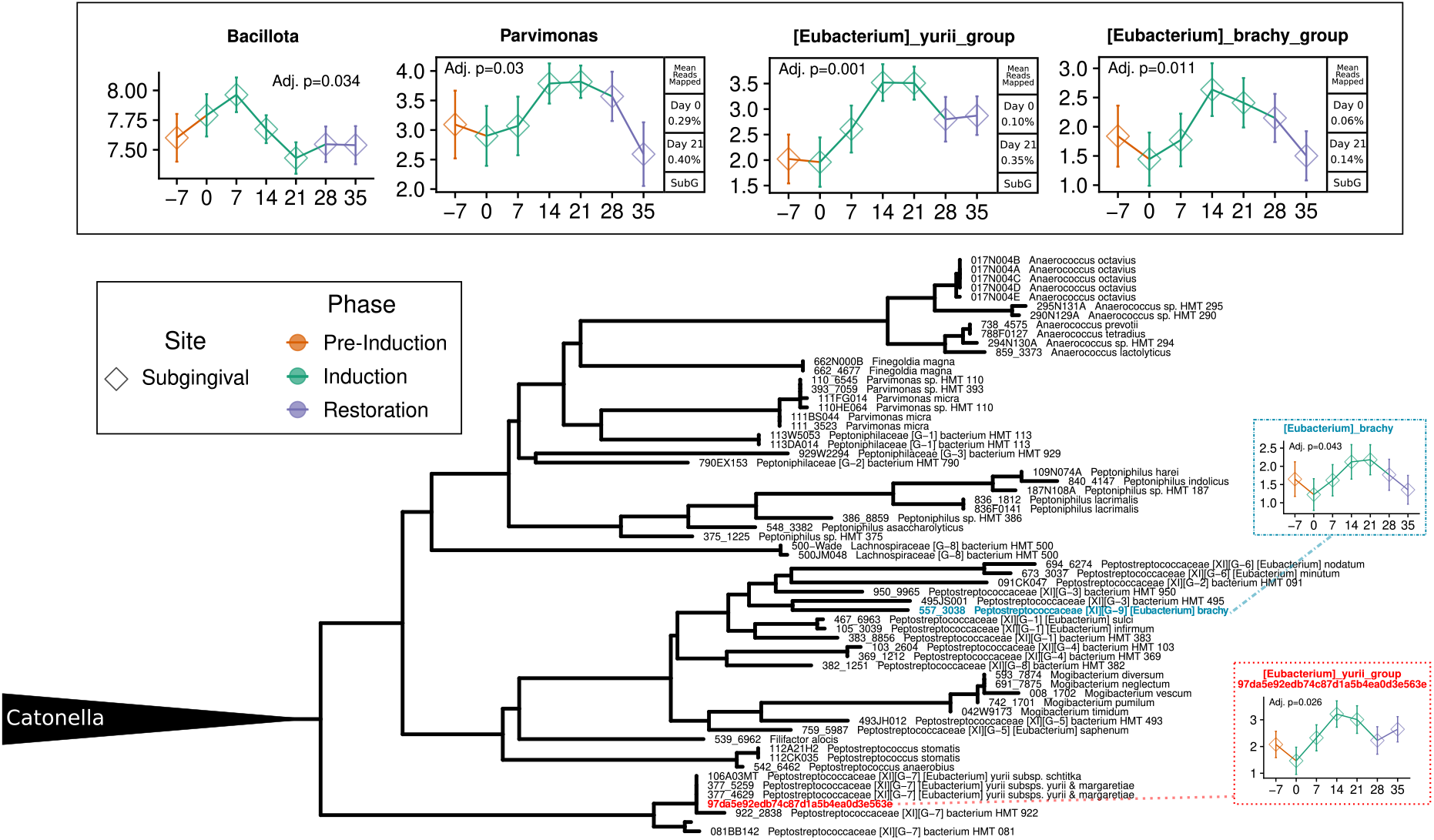
Parvimonas. Sequence abundances aggregated at the phylum and genus level (solid black outline), species level (blue dashed outline) and ASV level (red dotted outline). Y-axis is CLR-transformed reads and x-axis is days. Phylogenetic tree situates differentially abundant named species (blue) and ASVs (red) alongside the eHOMD reference sequences (black). p-values are the result of testing the null hypothesis that the slope is zero over the induction phase, with only significant results shown.

**Figure S13.**
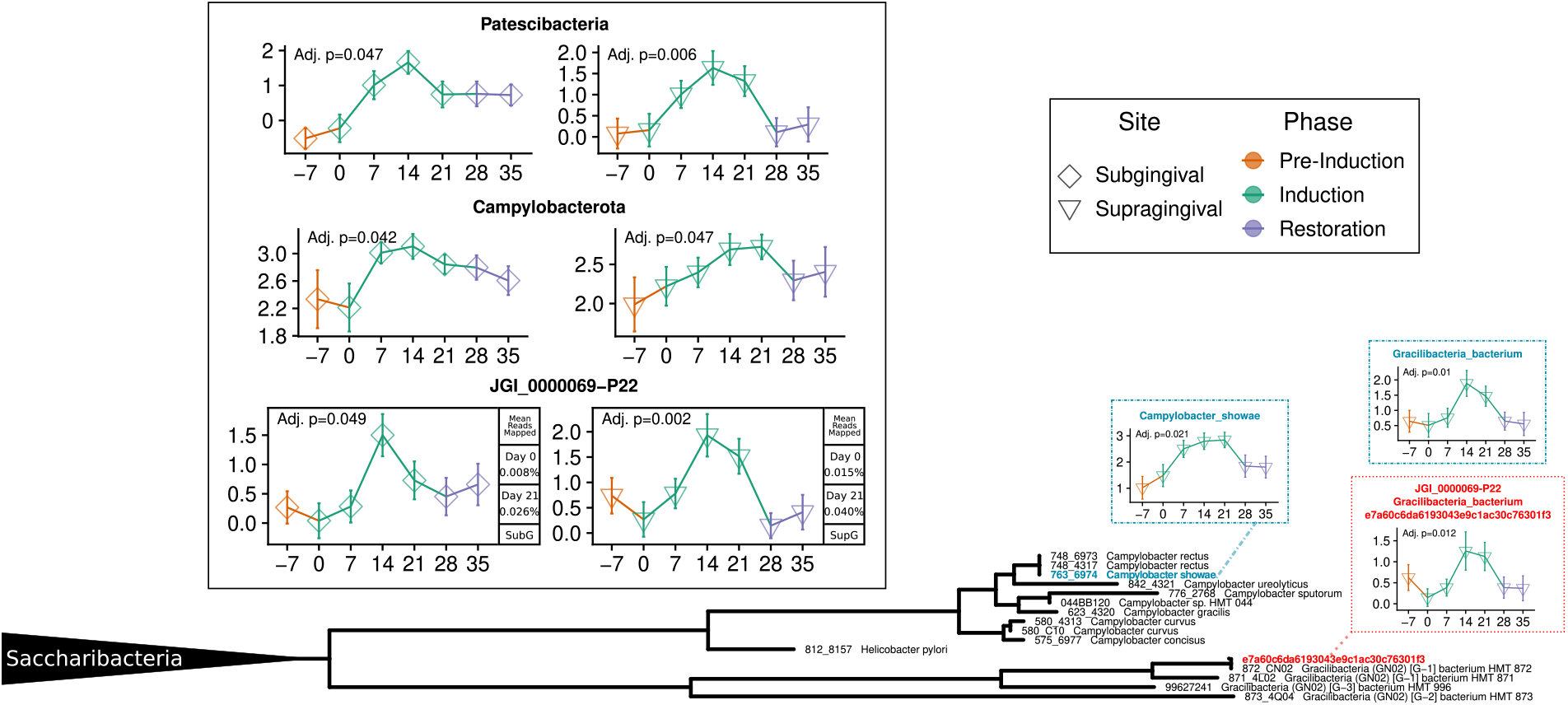
Campylobacter, Gracilibacteria (GN02, BD1-5, or SN-2) Sequence abundances aggregated at the phylum and genus level (solid black outline), species level (blue dashed outline) and ASV level (red dotted outline). Y-axis is CLR-transformed reads and x-axis is days. Phylogenetic tree situates differentially abundant named species (blue) and ASVs (red) alongside the eHOMD reference sequences (black). p-values are the result of testing the null hypothesis that the slope is zero over the induction phase, with only significant results shown.

**Figure S14.**
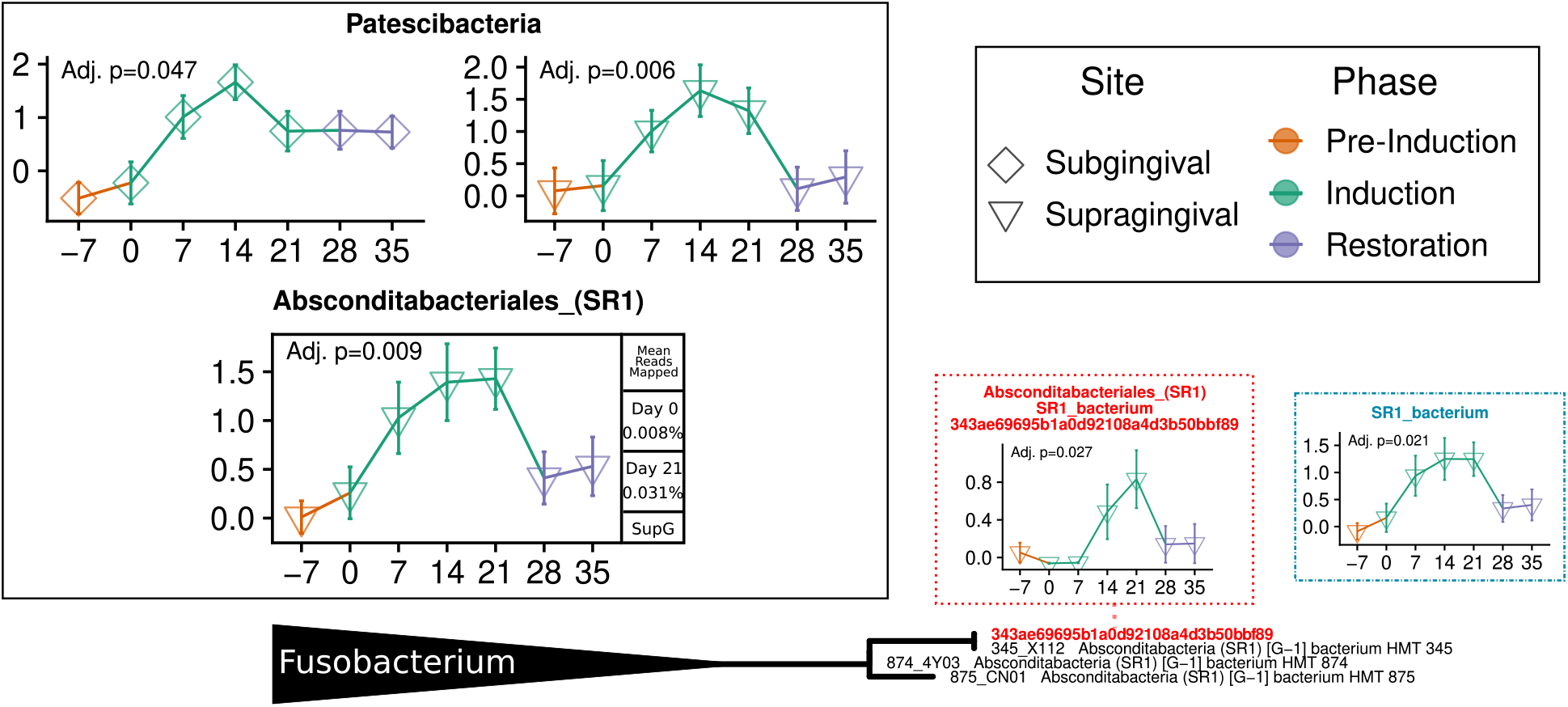
Absconditabacteriales. Sequence abundances aggregated at the phylum and genus level (solid black outline), species level (blue dashed outline) and ASV level (red dotted outline). Y-axis is CLR-transformed reads and x-axis is days. Phylogenetic tree situates differentially abundant named species (blue) and ASVs (red) alongside the eHOMD reference sequences (black). p-values are the result of testing the null hypothesis that the slope is zero over the induction phase, with only significant results shown.

**Figure S15.**
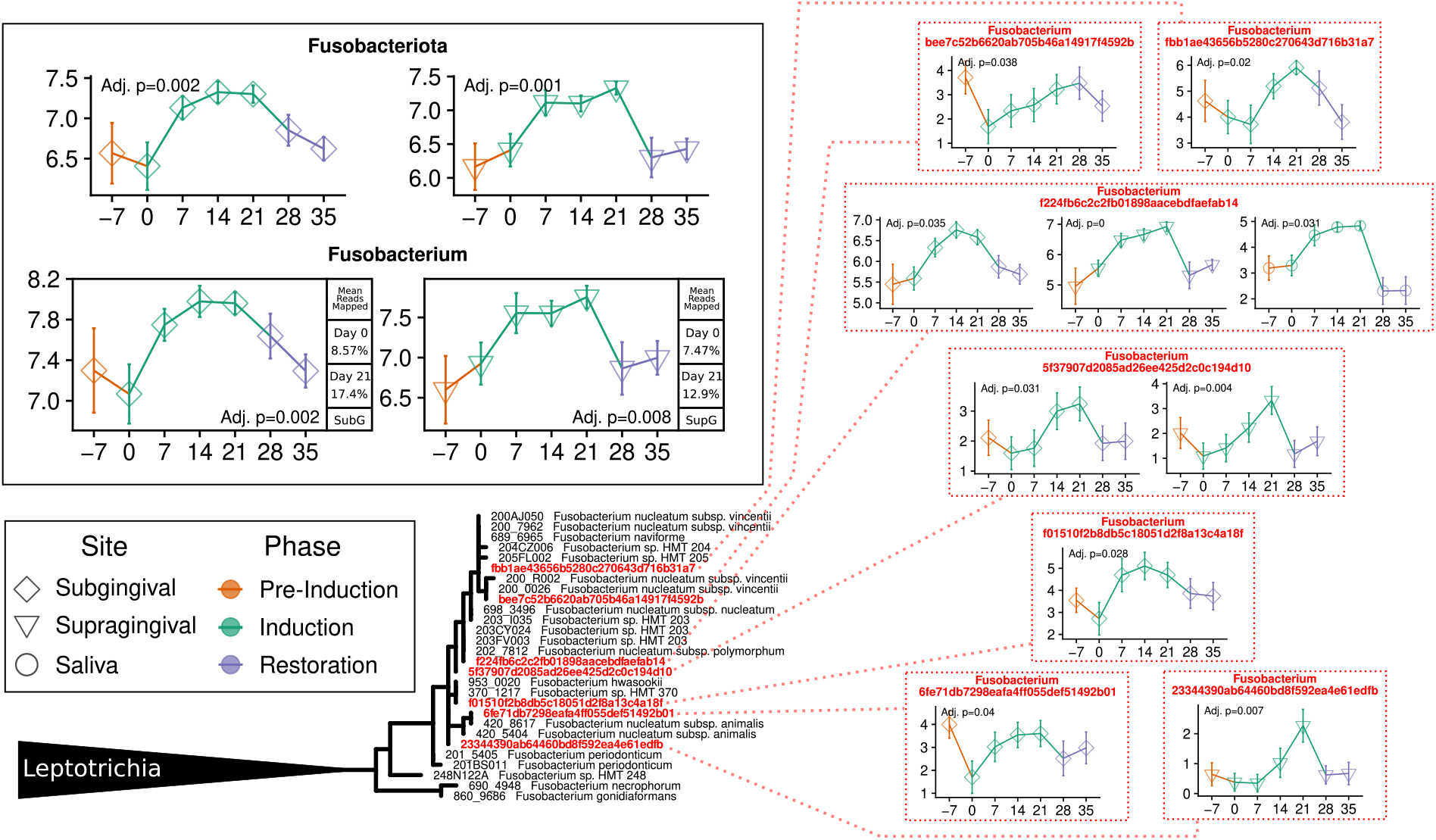
Fusobacteria. Sequence abundances aggregated at the phylum and genus level (solid black outline), species level (blue dashed outline) and ASV level (red dotted outline). Y-axis is CLR-transformed reads and x-axis is days. Phylogenetic tree situates differentially abundant named species (blue) and ASVs (red) alongside the eHOMD reference sequences (black). p-values are the result of testing the null hypothesis that the slope is zero over the induction phase, with only significant results shown.

**Figure S16.**
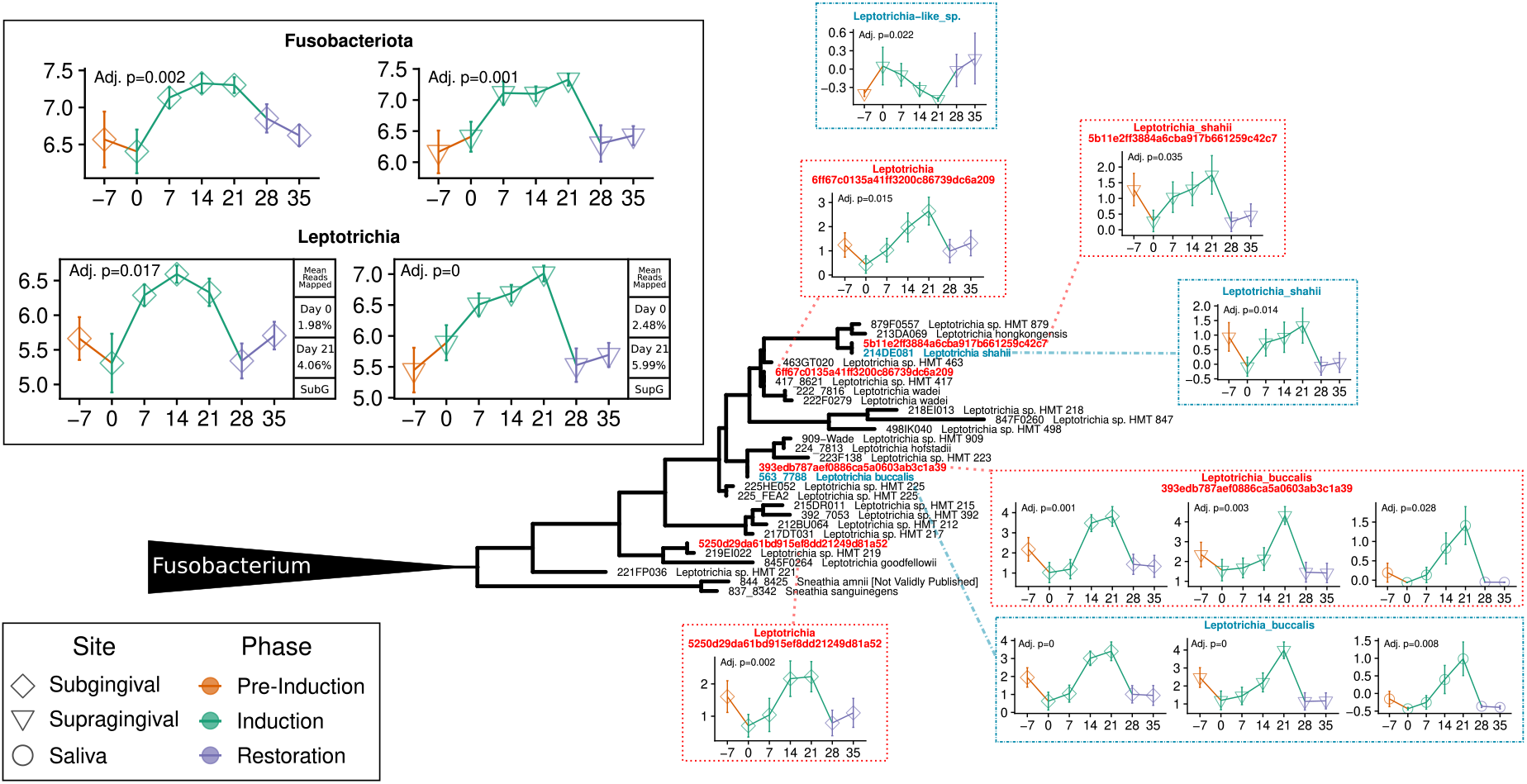
Leptotrichia. Sequence abundances aggregated at the phylum and genus level (solid black outline), species level (blue dashed outline) and ASV level (red dotted outline). Y-axis is CLR-transformed reads and x-axis is days. Phylogenetic tree situates differentially abundant named species (blue) and ASVs (red) alongside the eHOMD reference sequences (black). p-values are the result of testing the null hypothesis that the slope is zero over the induction phase, with only significant results shown.

**Figure S17.**
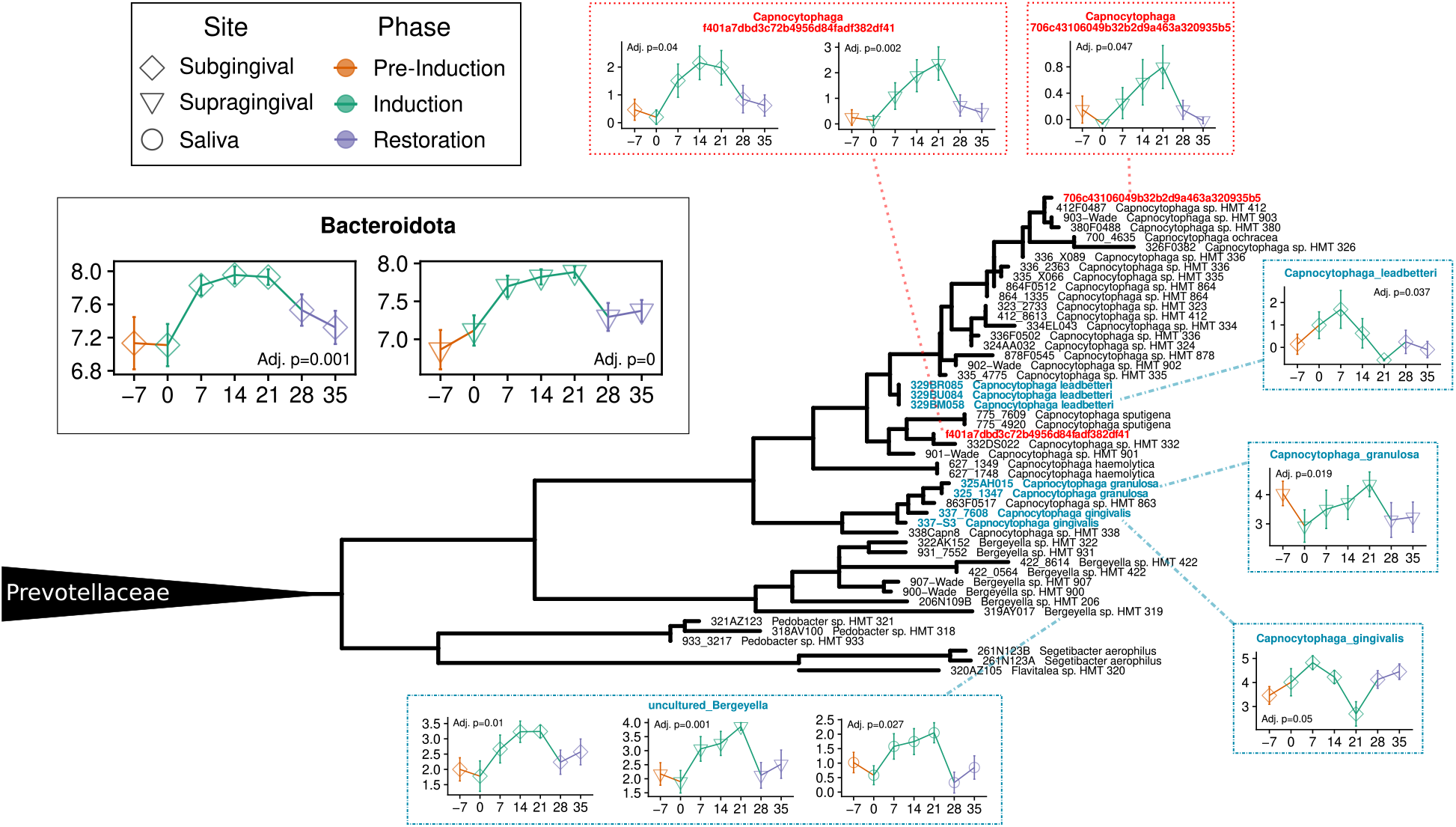
Capnocytophaga, Bergeyella. Sequence abundances aggregated at the phylum and genus level (solid black outline), species level (blue dashed outline) and ASV level (red dotted outline). Y-axis is CLR-transformed reads and x-axis is days. Phylogenetic tree situates differentially abundant named species (blue) and ASVs (red) alongside the eHOMD reference sequences (black). p-values are the result of testing the null hypothesis that the slope is zero over the induction phase, with only significant results shown.

**Figure S18.**
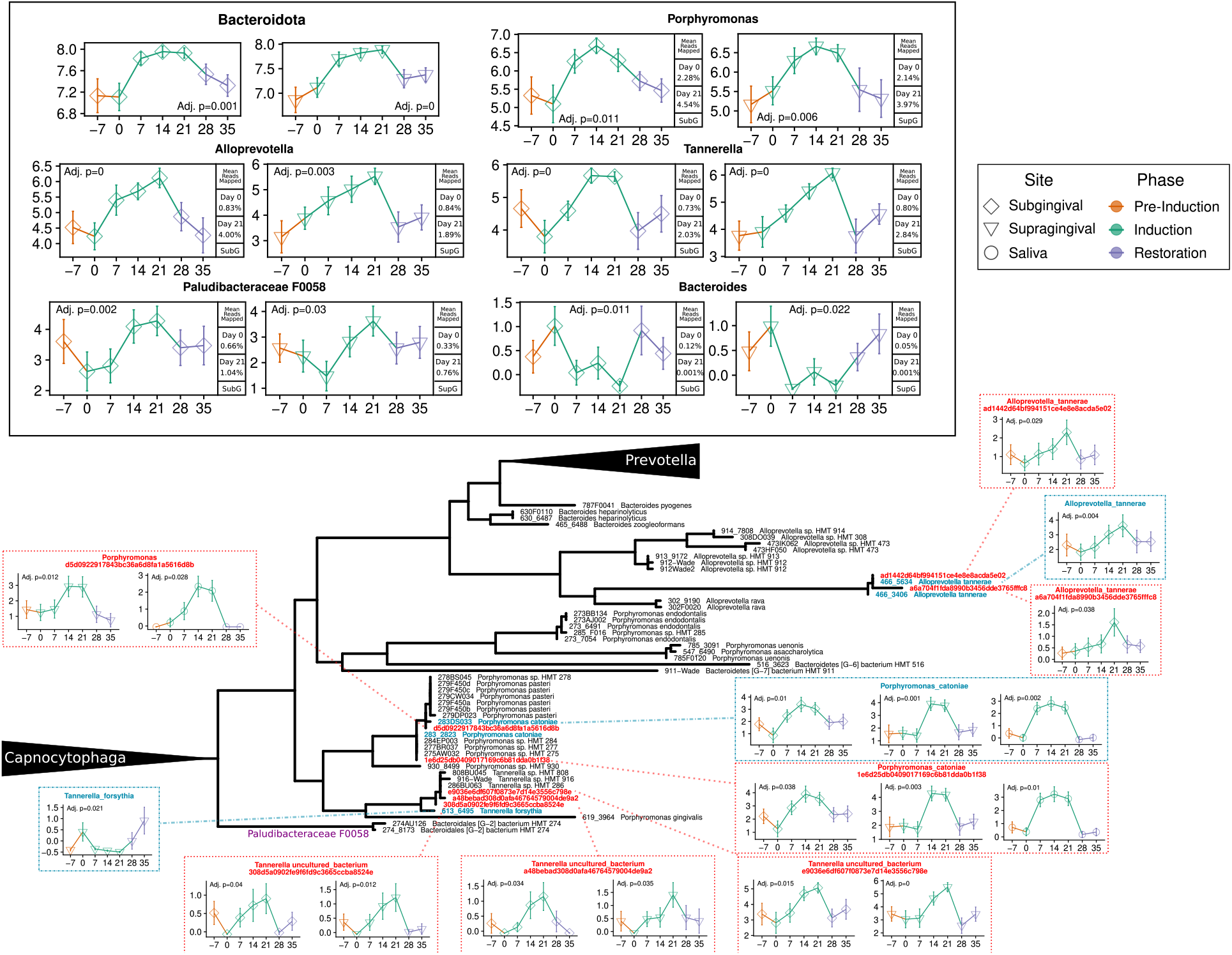
Alloprevotella, Tannerella, Porphyromonas. Sequence abundances aggregated at the phylum and genus level (solid black outline), species level (blue dashed outline) and ASV level (red dotted outline). Y-axis is CLR-transformed reads and x-axis is days. Phylogenetic tree situates differentially abundant named species (blue) and ASVs (red) alongside the eHOMD reference sequences (black). p-values are the result of testing the null hypothesis that the slope is zero over the induction phase, with only significant results shown.

**Figure S19.**
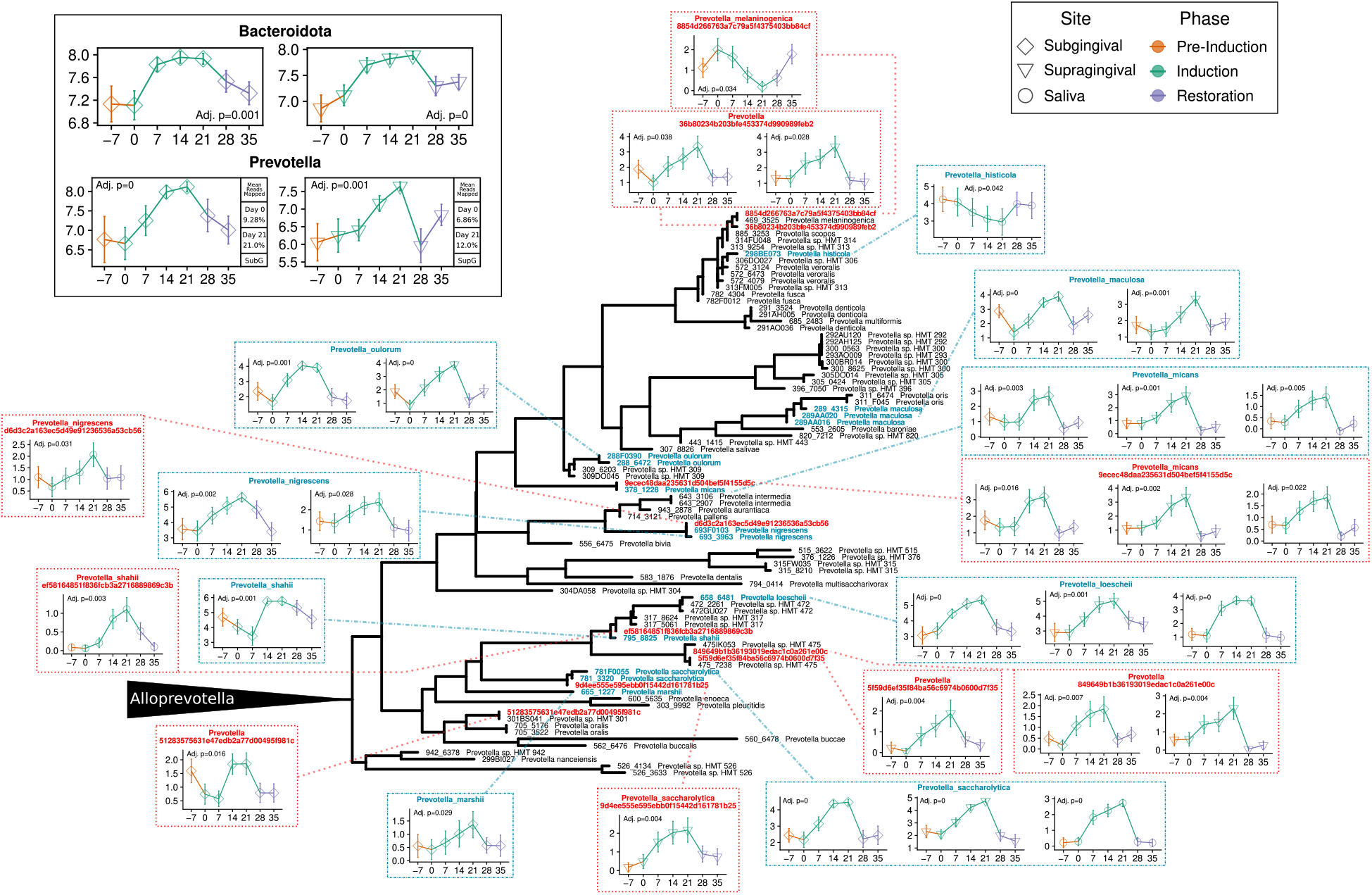
Prevotella. Sequence abundances aggregated at the phylum and genus level (solid black outline), species level (blue dashed outline) and ASV level (red dotted outline). Y-axis is CLR-transformed reads and x-axis is days. Phylogenetic tree situates differentially abundant named species (blue) and ASVs (red) alongside the eHOMD reference sequences (black). p-values are the result of testing the null hypothesis that the slope is zero over the induction phase, with only significant results shown.

**Figure S20.**
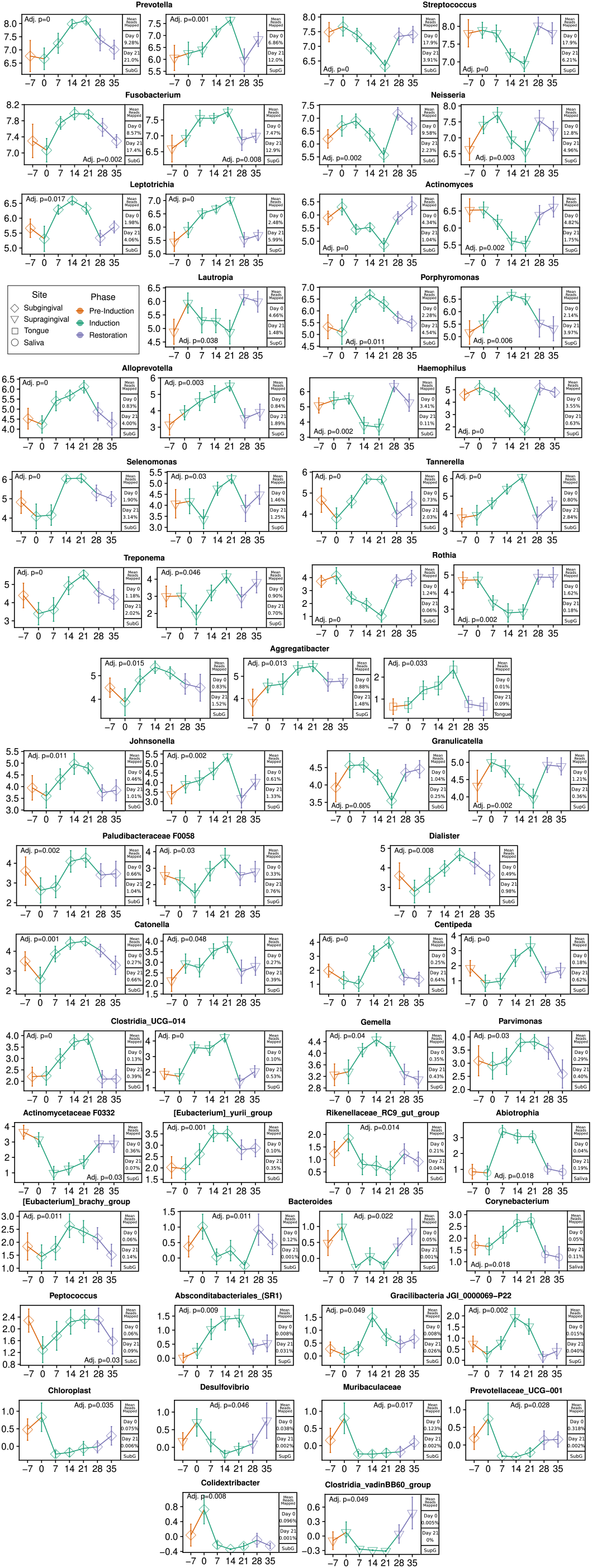
All genus-level designations with significantly increasing/decreasing abundance over the induction phase.

